# Amplification-free CRISPR/Cas13a-based viroid detection in RNA extracts from infected plants

**DOI:** 10.64898/2026.07.02.736049

**Authors:** Linh T.T. Le, Roser Montagud-Martínez, Guillermo Rodrigo, José-Antonio Daròs

## Abstract

Viroids are plant infectious agents that threaten agricultural production. Current viroid detection methods rely on RT-PCR–based assays, which require specialized laboratory equipment and can sometimes produce false-negative results or non-specific amplification due to the high sequence conservation among closely related viroid species. CRISPR-based diagnostics, particularly Cas12-based systems for DNA detection (DETECTR) and Cas13a-based systems (SHERLOCK) for RNA detection, have emerged as powerful tools for nucleic acid diagnostics. However, most existing workflows still rely on target amplification and, in the case of Cas13a systems, require additional *in vitro* transcription steps, limiting their simplicity and direct applicability for plant diagnostics. Here, we developed a direct amplification-free Cas13a-based detection platform for viroids using potato spindle tuber viroid (PSTVd) as a model. We optimized CRISPR RNA (crRNA) design, identified inhibitory effects of plant total RNA on readout signal, and employed simplified viroid RNA enrichment workflows enabling robust detection in plant samples. The system further supported both PSTVd-specific and broad-spectrum pospiviroid (genus *Pospiviroid*) detection and was successfully extended to avocado sunblotch viroid (family *Avsunviroidae*), demonstrating its adaptability across distinct viroid families. Together, these results establish a practical and modular Cas13a-based platform, not only for viroid diagnostics, but also for broader applications in RNA-derived plant pathogen detection.

**Graphical abstract:** 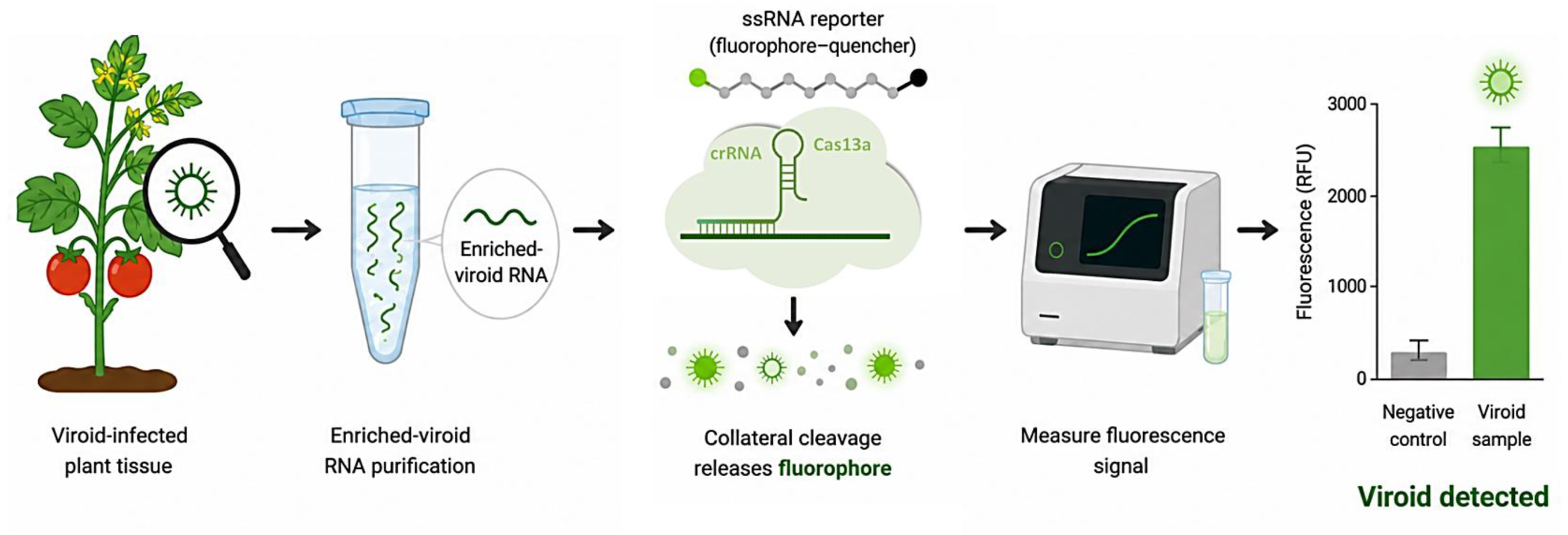

**Significance statement:** A simplified RNA enrichment workflow combined with CRISPR-Cas13a enables direct, amplification-free detection of plant viroids. The assay supports early and reliable diagnosis across different tomato varieties and provides a practical strategy for improving molecular detection of plant pathogens.

## Introduction

Viroids are the smallest known infectious agents infecting plants, consisting solely of small circular single-stranded RNAs ranging from 246 to 401 nucleotides (nt), approximately ten-fold smaller than the genomes of the smallest known RNA viruses (Hao et al., 2024; Kalantidis et al., 2025; Ortolá and Daròs, 2023; Sano, 2021). Viroid molecules adopt rod-like or branched secondary structures with a high degree of self-complementarity that promotes compact RNA folding (Pedrelli and Vergine, 2026; Sanger et al., 1976; Steger and Perreault, 2016). Despite their minimal genomes and lack of protein-coding capacity, viroids can cause severe diseases in numerous economically important crops, resulting in substantial agricultural and horticultural losses worldwide (Di Serio et al., 2023; Hammond, 2017; Venkataraman et al., 2024). Currently, 44 formally recognized viroid species are classified into two families: *Pospiviroidae*, comprising 39 members, and *Avsunviroidae*, comprising 5 members (Di Serio et al., 2021). Viroids belonging to the family *Pospiviroidae* are considered major quarantine pathogens in several countries due to their high transmissibility and economic impact on agricultural production (Verhoeven, 2010). Specifically, potato spindle tuber viroid (PSTVd; *Pospiviroid solani*) was the first viroid discovered and remains one of the most extensively studied viroid species (Diener, 1971). PSTVd represents an important phytosanitary concern because of its broad host range (Singh, 1973), efficient mechanical and seed transmission, and significant impact on tomato and potato production (Hussain et al., 2026).

Current viroid diagnostic methods mainly rely on nucleic acid-based approaches such as reverse transcription (RT)-polymerase chain reaction (PCR), RT-quantitative PCR (qPCR), RT-loop-mediated isothermal amplification (LAMP), hybridization assays, and next-generation sequencing (Nie and Singh, 2017; Pallás et al., 2018; Zhang and Li, 2024). Among these, RT-qPCR remains the most widely used approach because of its high sensitivity and specificity (Guček et al., 2023; Hajeri et al., 2022; Leichtfried et al., 2021). However, these techniques generally require specialized laboratory infrastructure, expensive instrumentation, and trained personnel. In addition, the high sequence conservation among closely related viroids makes the design of highly specific primers challenging, potentially affecting accurate discrimination between viroid species. Diagnostic strategies based on the clustered regularly interspaced short palindromic repeats (CRISPR) and CRISR-associated (Cas) nucleases from bacteria and archaea have recently emerged as powerful tools for nucleic acid detection. Many CRISPR/Cas-based nucleic acid biosensors have been developed to detect nucleic acids of viral and bacterial pathogens in clinical samples, as well as other applications in life sciences including biosecurity, food safety and environmental assessment. Additionally, CRISPR/Cas-based nucleic acid detection systems have demonstrated higher specificity compared with many conventional molecular diagnostic method (Tian et al., 2026; Zhou et al., 2024). In particular, the programmable recognition and cleavage activities of CRISPR/Cas proteins enable precise targeting of specific nucleic acid sequences, making these systems powerful tools for sensitive and accurate nucleic acid diagnostics (Gong et al., 2025; Jaybhaye et al., 2024; Montagud-Martínez and Márquez-Costa, 2025). In particular, Cas12-based systems such as the DNA endonuclease targeted CRISPR trans reporter (DETECTR) have been widely applied for DNA detection, whereas Cas13a-based platforms such as the specific high-sensitivity enzymatic reporter un-locking (SHERLOCK) enable RNA detection through target-activated non-specific collateral cleavage of fluorescent RNA reporters (Chertow, 2018; Gootenberg et al., 2017; Li et al., 2018). Since the development of the SHERLOCK platform, Cas13a-based diagnostics have been successfully applied for the detection of numerous human, animal, and plant pathogens, including viroids (Anwar et al., 2025; Khan et al., 2021; Yao et al., 2021; Zhai et al., 2024). However, most currently available CRISPR diagnostic workflows still depend on nucleic acid amplification to achieve sufficient analytical sensitivity. Furthermore, Cas13a-based assays commonly require additional *in vitro* transcription steps before detection, increasing workflow complexity, assay time, contamination risk, and limiting direct applicability for plant diagnostics.

Recent studies have demonstrated the feasibility of DETECTR and SHERLOCK detection for viroids using amplification-assisted workflows (Jiao et al., 2021) (Zhai et al., 2024). Nevertheless, direct amplification-free Cas13a detection of viroids from plant-derived RNA samples has not yet been established. In particular, the effects of complex plant RNA backgrounds on Cas13a activity and the development of simplified RNA enrichment strategies compatible with direct Cas13a detection remain poorly understood. Addressing these limitations is essential for developing practical CRISPR-based diagnostic systems suitable for routine plant pathogen surveillance and field applications.

In this study, we established a direct amplification-free Cas13a-based detection strategy for viroids using PSTVd as a representative model. We approached CRISPR RNA (crRNA) selection and examined the impact of plant-derived total RNA on Cas13a detection performance, which guided the development of simplified viroid RNA enrichment procedures for reliable detection in plant tissues. To improve detection versatility, we further designed a dual-crRNA approach capable of both specific PSTVd detection and broader recognition of pospiviroids. The applicability of the platform was additionally evaluated using avocado sunblotch viroid (ASBVd; *Avsunviroid perseae*) belonging to a different viroid family (*Avsunviroidae*), demonstrating its adaptability to phylogenetically distinct viroids. Overall, this work provides a foundation for practical, modular, and amplification-free Cas13a-based diagnostics for plant viroids.

## Results

### Rational design and screening of crRNAs for Cas13a-based amplification-free detection of PSTVd

To establish an efficient amplification-free Cas13a-based detection platform for PSTVd, several crRNAs targeting distinct regions of the PSTVd genome were designed based on the predicted secondary structure of the viroid RNA (Figure 1A). Candidate target sites were distributed across several domains of PSTVd, including the terminal left (TL), terminal conserved region (TCR), pathogenicity (P), central (C), central conserved region (CCR), and variable (V) domains, to evaluate the influence of target accessibility and local RNA structure on Cas13a activation efficiency.

**Figure 1.**
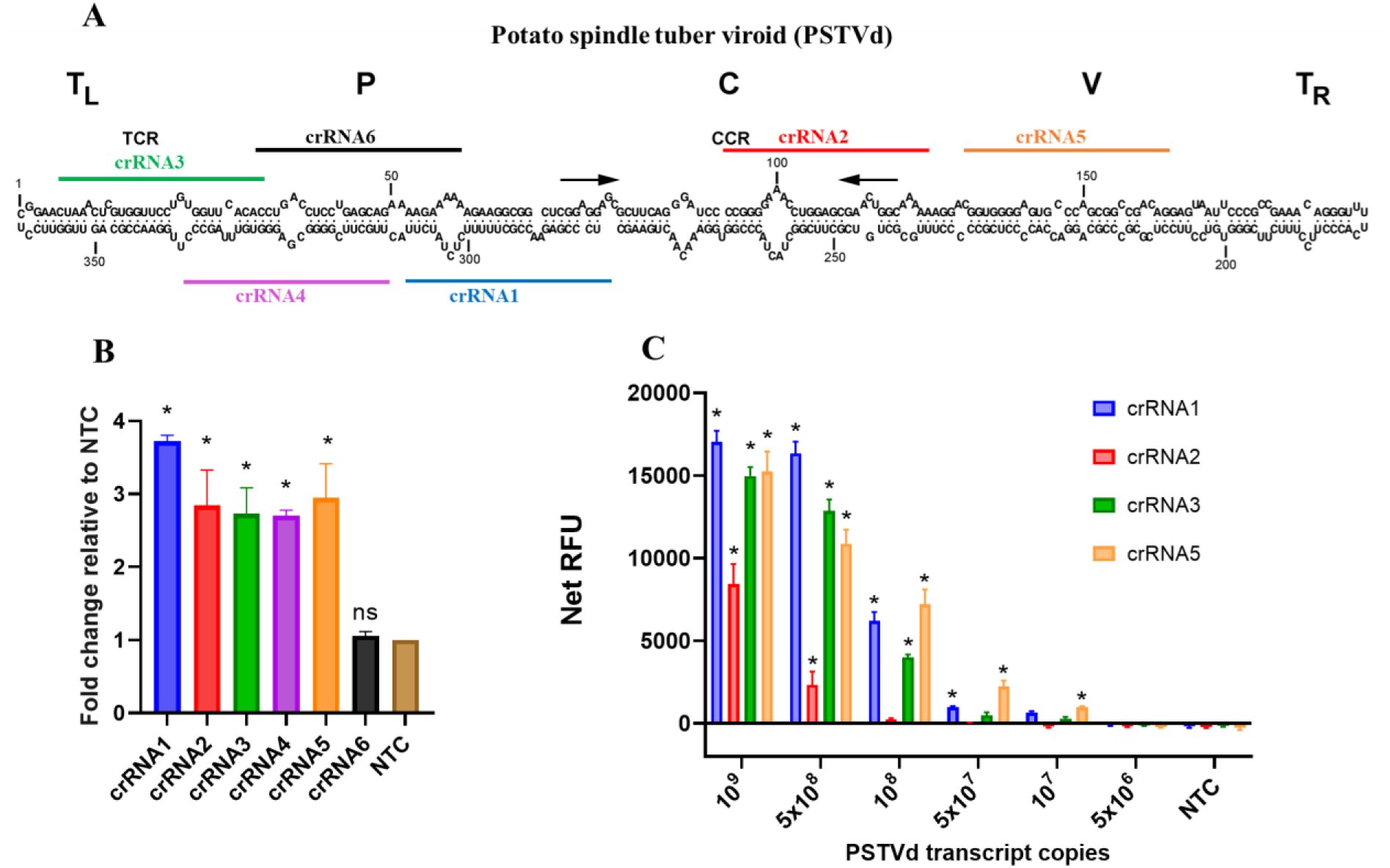
Screening and sensitivity evaluation of crRNAs targeting PSTVd for Cas13a-based detection. (A) Schematic representation of the predicted secondary structure of PSTVd and the positions of the designed crRNA target sites across different domains. (B) Initial screening of candidate crRNAs using *in vitro*-transcribed PSTVd RNA in a fluorimetric assay. Fluorescence signals are presented as fold change relative to the no-target control (NTC). (C) Sensitivity evaluation of selected crRNAs using serial dilutions of *in vitro*-transcribed PSTVd RNA. Net relative fluorescence units (Net RFU) are plotted versus PSTVd transcript copies. (B and C) *, statistically significant *p* = 0.001; ns, non-significant.

The detection performance of each crRNA was initially evaluated using *in vitro*-transcribed PSTVd RNA in a fluorescence assay based on the *trans*-cleavage activity of Cas13a on a 6-nt-long RNA probe labelled at 5’ and 3’ ends, respectively, with fluorescence and the quencher Iowa Black. Among the six tested candidates, crRNA1 and crRNA5 generated the strongest fluorescence signals, exhibiting approximately 3.5–3.8-fold higher fluorescence relative to the control with no viroid transcript added. In contrast, crRNA6 failed to produce significant signal enhancement compared with the no-viroid control (Figure 1B). Based on the initial screening results, the four best-performing crRNAs (crRNA1, crRNA2, crRNA3, and crRNA5) were selected for further sensitivity evaluation using serial dilutions of *in vitro*-transcribed PSTVd RNA (Figure 1C). At high transcript concentrations (10⁹–5 × 10⁸ copies), all selected crRNAs produced fluorescence signals significantly above the no-viroid control. However, fluorescence intensity progressively decreased with lower target copy numbers, and differences in detection performance became more pronounced. Among the tested candidates, crRNA1 generated the highest fluorescence intensity at high transcript concentrations, whereas crRNA5 retained detectable fluorescence signals at lower target concentrations. The lowest detectable concentration observed for crRNA5 was 10⁷ transcript copies (∼16.6 pM), followed by 5 × 10⁷ copies for crRNA1 under the tested conditions. These results indicate that each crRNA exhibited distinct analytical sensitivity profiles, with crRNA5 demonstrating the best apparent sensitivity among the tested candidates. Therefore, crRNA5 was selected for subsequent assay optimization and downstream experiments.

### Plant total RNA interferes with direct Cas13a-based viroid detection

After establishing the optimized Cas13a detection system using *in vitro*-transcribed PSTVd RNA, we next evaluated its applicability for direct detection in plant-derived RNA samples. RNA extracted from PSTVd-infected tomato plants was initially processed using a previously reported viroid enrichment workflow based on phenol extraction followed by CF11 cellulose chromatography. Although this method has been widely used for efficient viroid RNA enrichment, unexpectedly low Cas13a fluorescence signals were observed when the enriched plant RNA samples were directly subjected to the detection assay. To further investigate this phenomenon, Cas13a fluorescence signals were compared among non-infected, infected, non-infected plus PSTVd transcripts, infected plus PSTVd transcripts, PSTVd transcripts-only, and no-target control reactions (Figure 2A). Both infected and non-infected plant RNA samples alone generated only weak fluorescence signals that were not significantly different from the no-target control. In contrast, substantially stronger fluorescence signals were observed in reactions containing *in vitro*-transcribed PSTVd target RNA. Notably, the *in vitro* transcribed PSTVd-only reaction produced higher fluorescence intensity than reactions containing *in vitro* PSTVd transcripts supplemented with infected or non-infected plant RNA extracts. This finding suggests that components of the plant-derived RNA preparations interfere with efficient Cas13a activation when present at high concentrations.

**Figure 2.**
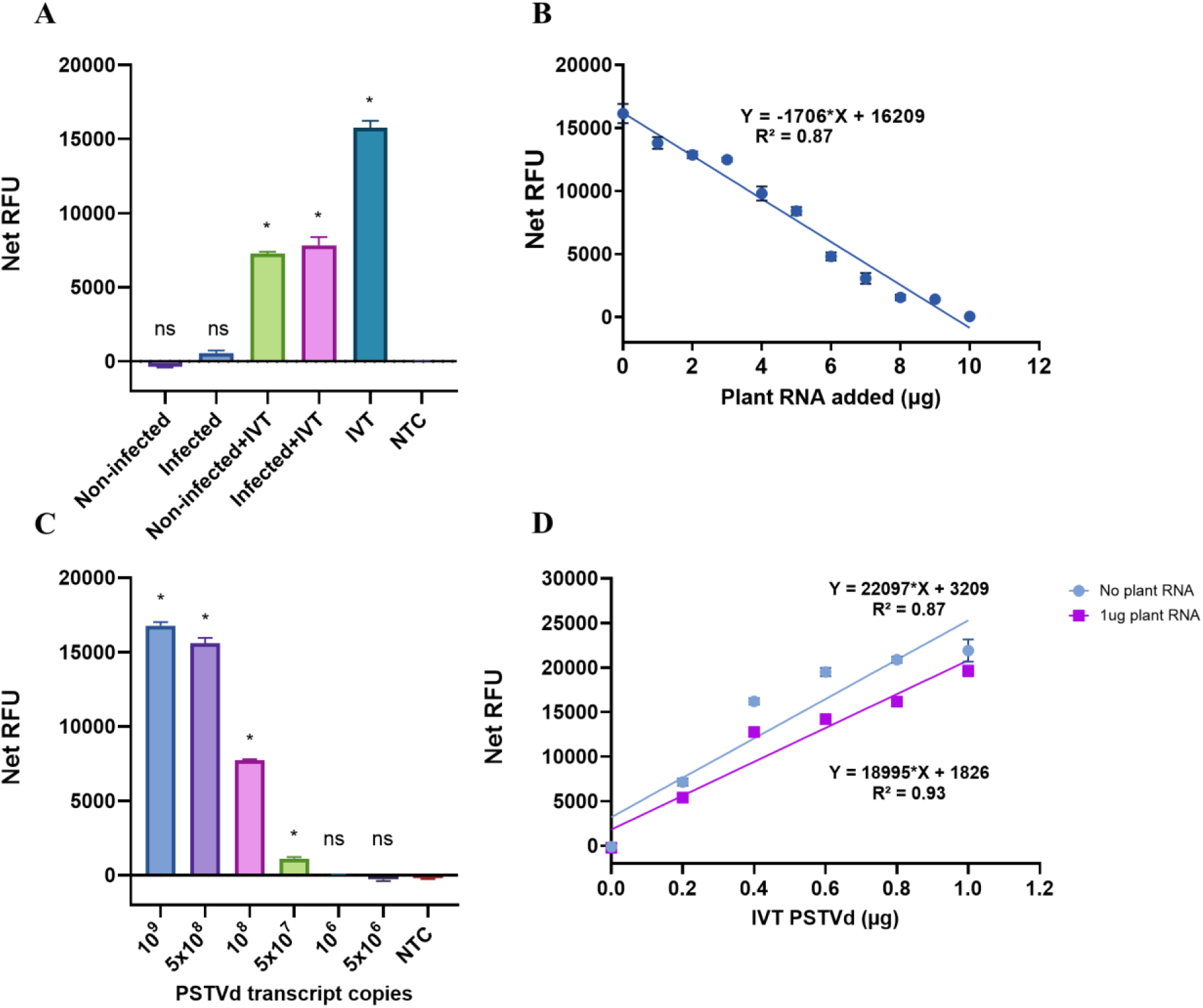
Analysis of total RNA interference on Cas13a-based PSTVd detection. (A) Net RFU from non-infected, infected, non-infected plus *in vitro* transcribed (IVT), infected plus IVT, IVT-only, and no-target control (NTC) reactions. (B) Effect of increasing amounts of total plant RNA on Cas13a fluorescence output in reactions containing 1 ng IVT target RNA. (C) Effect of adding fixed 1µg plant-derived background RNA on the limit of detection of the Cas13a assay in reactions containing 1ng IVT target RNA. (D) Cas13a fluorescence responses to increasing IVT concentrations (0–1 ng) in the presence or absence of 1 µg total plant background RNA. (A and C) *, statistically significant *p* = 0.001; ns, non-significant.

We next investigated the effect of plant-derived background RNA on Cas13a detection performance by adding increasing amounts of total RNA extracted from non-infected plants into reactions containing a fixed amount (1 ng) of *in vitro*-transcribed PSTVd RNA. Fluorescence output progressively decreased as the amount of total RNA increased (Figure 2B), and linear regression analysis revealed a strong negative correlation between background RNA concentration and fluorescence intensity. Notably, inhibitory effects were still observed after additional silica column purification or DNase treatment of the plant heterologous RNA, suggesting that inhibition is primarily associated with the abundance of plant-derived background RNA rather than residual extraction contaminants or DNA.

To further evaluate assay sensitivity under conditions more representative of plant samples, serial dilutions of *in vitro*-transcribed PSTVd RNA were tested in the presence of 1µg total RNA from non-infected plants (Figure 2C). Under these background RNA conditions, the detection limit shifted from 10⁷ copies observed with *in vitro*-transcribed PSTVd RNA alone (Figure 1C) to approximately 5 × 10⁷ copies, supporting that plant-derived RNA backgrounds reduce Cas13a detection sensitivity. Finally, to determine whether low levels of background RNA could still be tolerated, increasing concentrations of *in vitro*-transcribed PSTVd RNA were tested in the presence or absence of 1 µg plant total RNA (Figure 2D). In both conditions, fluorescence signals increased proportionally with increasing target RNA concentration, and linear regression analysis showed comparable fluorescence response trends between the two conditions. Together, these findings indicate that while high concentrations of plant-derived RNA strongly inhibit Cas13a activity, low levels of background RNA can still be tolerated without substantially compromising detection performance.

### Selective LiCl-mediated RNA enrichment restores Cas13a detection sensitivity

Based on the inhibitory effects observed from plant-derived RNA backgrounds, we next evaluated RNA extraction and enrichment workflows designed to improve direct Cas13a-based PSTVd detection (Figure 3A). To reduce abundant host RNAs while retaining viroid-associated RNAs, a selective LiCl precipitation step was incorporated into the extraction workflow. Because low-molecular-weight RNAs such as viroids preferentially remain soluble in LiCl, whereas a large proportion of high-molecular-weight cellular RNAs are precipitated, this approach enabled selective reduction of background plant RNAs. Polyacrylamide gel electrophoresis (PAGE) analysis and reverse transcription (RT)-quantitative polymerase chain reaction (qPCR) confirmed successful PSTVd infection in all inoculated plants (Supplementary Figure S1). Comparison of the different workflows showed that incorporation of the LiCl enrichment step substantially improved Cas13a fluorescence signals in PSTVd-infected samples (Figure 3B). Among the tested methods, the complete phenol/CF11/LiCl workflow generated the strongest fluorescence output. In contrast, silica column-based RNA purification produced markedly weaker fluorescence signals, consistent with limited removal of inhibitory background RNAs and lack of viroid enrichment. Importantly, robust fluorescence signals were still obtained using the simplified phenol/LiCl workflow without the CF11 chromatography step (Figure 3B). Although fluorescence intensity was moderately reduced compared with the complete phenol/CF11/LiCl workflow, infected samples remained clearly distinguishable from controls and generated substantially higher fluorescence signals than silica column-purified RNA preparations. Real-time fluorescence kinetics further demonstrated rapid Cas13a activation using both LiCl-based workflows, with strong fluorescence accumulation observed within the first 10 min of the reaction (Figure 3C). In contrast, silica column-purified RNA produced substantially slower fluorescence kinetics and markedly reduced fluorescence intensity. Together, these results indicate that reducing background plant RNAs substantially improves Cas13a detection performance in plant-derived samples.

**Figure 3.**
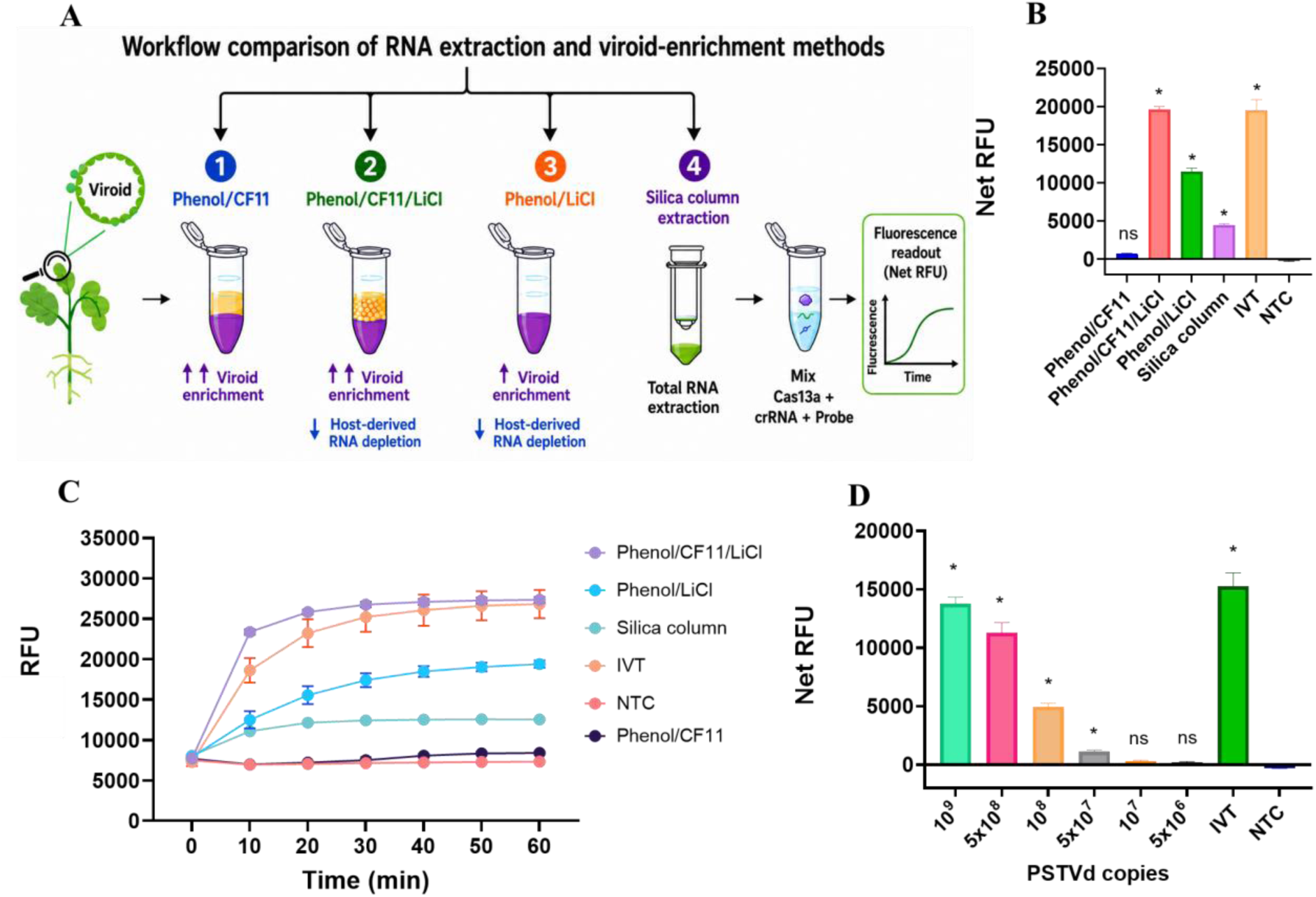
LiCl-based RNA enrichment improves Cas13a-mediated PSTVd detection. (A) Schematic overview of the RNA extraction and enrichment workflows evaluated for Cas13a-based PSTVd detection, including phenol/CF11, phenol/CF11/LiCl, phenol/LiCl, and silica column-based extraction methods. (B) Net RFU comparison of different RNA extraction and enrichment workflows. (C) Real-time Cas13a fluorescence kinetics generated from different RNA extraction and enrichment workflows. (D) Sensitivity evaluation of direct Cas13a-based PSTVd detection using serial dilutions of RNA extracted from PSTVd-infected tomato samples following the optimized phenol/LiCl enrichment workflow. (B and D) *, statistically significant *p* = 0.001; ns, non-significant.

To further evaluate the applicability of the optimized workflow in biological samples, the analytical sensitivity of direct Cas13a-based PSTVd detection was assessed using serial dilutions of RNA extracted from infected tomato plants following the simplified phenol/LiCl workflow (Figure 3D). PSTVd copy numbers in plant-derived RNA samples were estimated using absolute RT-qPCR prior to Cas13a analysis (Supplementary Data 1). The limit of detection of the optimized amplification-free Cas13a assay was estimated to be approximately 5 × 10⁷ PSTVd copies, corresponding to ∼83 pM.

Overall, although the complete phenol/CF11/LiCl workflow produced the strongest fluorescence signals, the simplified phenol/LiCl workflow maintained robust detection performance while substantially reducing workflow complexity and processing time. Therefore, the simplified workflow was selected for subsequent plant-sample detection experiments.

### Robust and early Cas13a-based viroid detection across multiple tomato varieties

To evaluate the robustness and broad applicability of the optimized Cas13a detection workflow in plant samples, four tomato varieties with distinct PSTVd accumulation profiles were analyzed under identical experimental conditions, including inoculation procedure, growth environment, sampling time, and RNA preparation, phenol/LiCl workflow. Tomato plants from the Rutgers, Micro-Tom, Moneymaker, and Marglobe cultivars were agroinoculated with PSTVd, while corresponding non-infected plants were included as negative controls. PAGE analysis and RT-qPCR confirmed successful PSTVd infection in all inoculated plants and revealed clear differences in viroid accumulation among the tested tomato varieties (Supplementary Figure S2). The Cas13a assay successfully detected PSTVd in all infected samples (Figure 4A). These results demonstrate that the simplified enrichment workflow combined with Cas13a detection is robust across diverse tomato genetic backgrounds.

**Figure 4.**
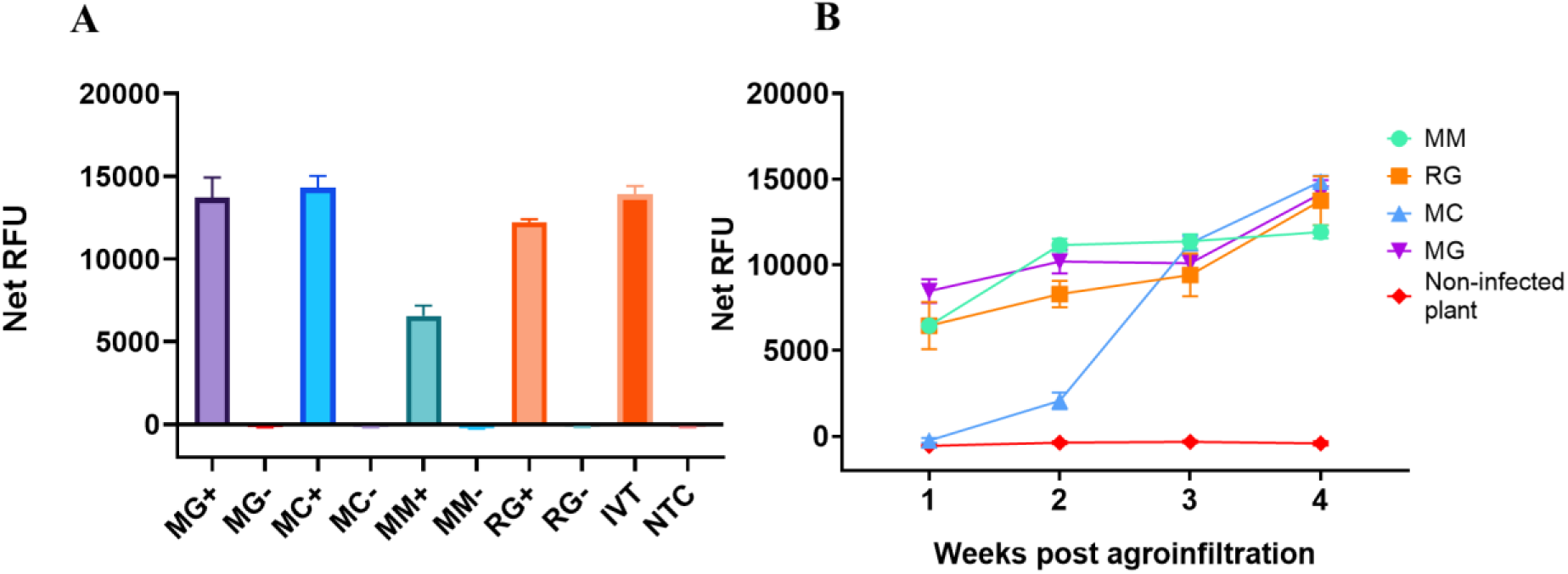
Robust and early Cas13a-based detection of PSTVd across multiple tomato varieties. (A) Net RFU obtained from PSTVd-infected and non-infected Rutgers (RG), Micro-Tom (MT), Moneymaker (MM), and Marglobe (MG) tomato plants using the optimized Cas13a detection workflow. (B) Time-course analysis of Cas13a-based PSTVd detection in four tomato varieties at 1, 2, 3, and 4 weeks post agroinfiltration (wpi). Fluorescence signals increased progressively throughout infection, reaching the highest levels at 4 wpi.

Among the tested varieties in our experimental conditions, Moneymaker exhibited the lowest viroid accumulation level based on RT–qPCR analysis, which corresponded to substantially lower fluorescence intensity in the Cas13a assay compared with the other infected tomato varieties (Figure 4A). In contrast, Rutgers, Micro-Tom, and Marglobe plants generated markedly stronger fluorescence signals, consistent with their higher PSTVd accumulation levels. In addition, silica column-purified RNA extracted from the four tomato varieties showed variable Cas13a detection efficiencies, and PSTVd could not be reliably detected in the Moneymaker variety (Supplementary Figure S3).

To further investigate the potential of the assay for early detection, leaf samples from four tomato varieties were collected at 1, 2, 3, and 4 weeks post agroinfiltration (wpi) with PSTVd and analyzed using the optimized Cas13a detection workflow. The assay successfully detected PSTVd throughout the course of infection in all four tomato varieties. Significant fluorescence signals relative to the non-infected control were observed as early as 1 wpi in Rutgers, Moneymaker, and Marglobe plants, whereas Micro-Tom reached statistical significance from 2 wpi onward (Supplementary Figure S4). Fluorescence signals increased progressively over time, reaching their highest levels at 4 wpi, consistent with the progressive accumulation of PSTVd during infection. In addition, Cas13a fluorescence exhibited a strong inverse correlation with RT-qPCR Cq values, indicating that increased Cas13a fluorescence corresponded to lower RT-qPCR Cq values and, consequently, higher PSTVd accumulation (Supplementary Figure S5). These findings demonstrate that this optimized Cas13a detection workflow enables early, sensitive, and reliable detection of PSTVd across diverse tomato genetic backgrounds while maintaining robust performance throughout disease progression.

### Specific detection of PSTVd and universal detection of related pospiviroids

To evaluate the specificity and broader applicability of the Cas13a detection platform, five selected crRNAs (crRNA1, crRNA2, crRNA3, crRNA4, and crRNA5) were tested against *in vitro*-transcribed RNAs representing PSTVd and five additional members of the genus *Pospiviroid*, namely citrus exocortis viroid (CEVd), chrysanthemum stunt viroid (CSVd), tomato chlorotic dwarf viroid (TCDVd), tomato apical stunt viroid (TASVd) and portulaca lantent viroid (PLVd).

Distinct detection patterns were observed among the tested crRNAs. crRNA5 exhibited high specificity toward PSTVd, generating strong fluorescence signals exclusively for this viroid species while showing negligible activity against the other tested pospiviroids. In contrast, crRNA3 displayed broad detection capability and successfully detected all tested pospiviroids with substantially elevated fluorescence signals compared with the negative control, indicating targeting of highly conserved sequence regions shared among members of the genus *Pospiviroid*. crRNA1 and crRNA4 showed intermediate specificity patterns, detecting PSTVd, TCDVd, and TASVd, whereas crRNA2 displayed broader but incomplete cross-reactivity by detecting PSTVd, CEVd, TCDVd, TASVd, and PLVd. These differences likely reflect sequence conservation and mismatch tolerance within the corresponding target regions. Based on these findings, a dual-crRNA Cas13a detection strategy was proposed in which crRNA3 is used for broad-spectrum pospiviroid screening, while crRNA5 serves as a PSTVd-specific identifier. This combined approach enables both universal detection and species-specific discrimination within a single Cas13a-based diagnostic platform.

### Extension of the Cas13a detection platform to the family *Avsunviroidae*

To further evaluate the versatility and adaptability of the developed Cas13a detection platform, the system was extended to detect ASBVd, the type member of the family *Avsunviroidae* that is evolutionarily distinct from members of the genus *Pospiviroid* (family *Pospiviroidae*). Six crRNAs targeting different regions of the ASBVd genome were designed and screened using *in vitro*–transcribed ASBVd RNA (Figure 6A). All six crRNAs generated fluorescence signals above the no-target control, although distinct differences in fluorescence intensity were observed among the tested crRNAs, indicating variable target accessibility and Cas13a activation efficiencies across different ASBVd genomic regions. Among the tested candidates, crRNA5 consistently produced the strongest fluorescence signal and was therefore selected for subsequent sensitivity and plant-sample detection experiments. The analytical sensitivity of the optimized ASBVd detection system was subsequently evaluated using serial dilutions of *in vitro*-transcribed ASBVd RNA (Supplementary Figure S6). The Cas13a assay successfully detected as low as 10^7^ ASBVd transcript copies while maintaining clear discrimination from negative controls, demonstrating analytical sensitivity comparable to that observed for PSTVd detection.

**Figure 5.**
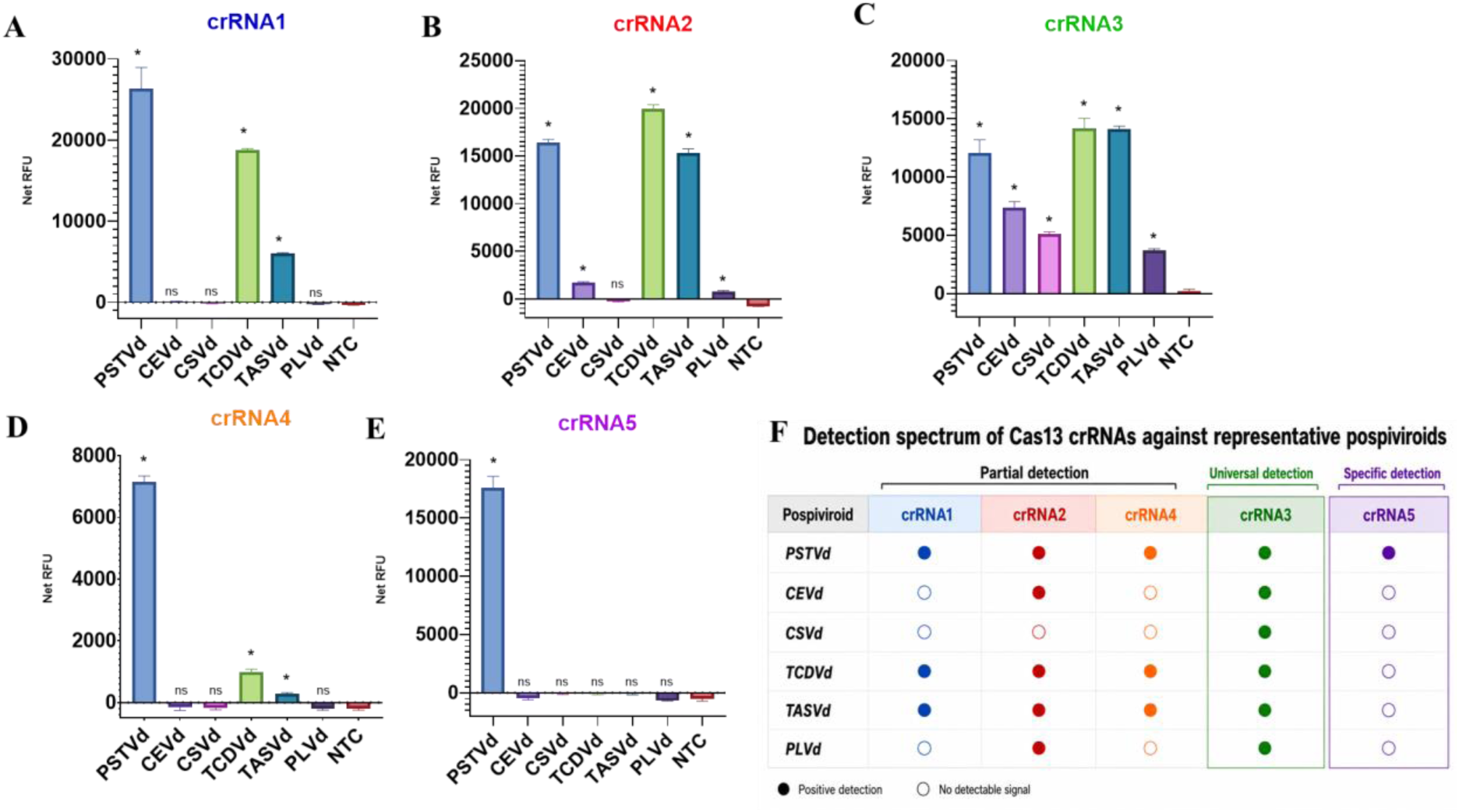
Specific and universal detection profiles of Cas13a crRNAs against representative pospiviroids. (A–E) Fluorescence-based detection of representative pospiviroids using Cas13a assays programmed with crRNA1 (A), crRNA2 (B), crRNA3 (C), crRNA4 (D), and crRNA5 (E). *In vitro*-transcribed RNAs corresponding to PSTVd, CEVd, CSVd, TCDVd, TASVd, and PLVd were tested individually. (F) Summary of detection spectra of the tested crRNAs against representative pospiviroids. Filled circles indicate positive detection, whereas open circles indicate no detectable signal. (A to E) *, statistically significant *p* = 0.001; ns, non-significant.

**Figure 6.**
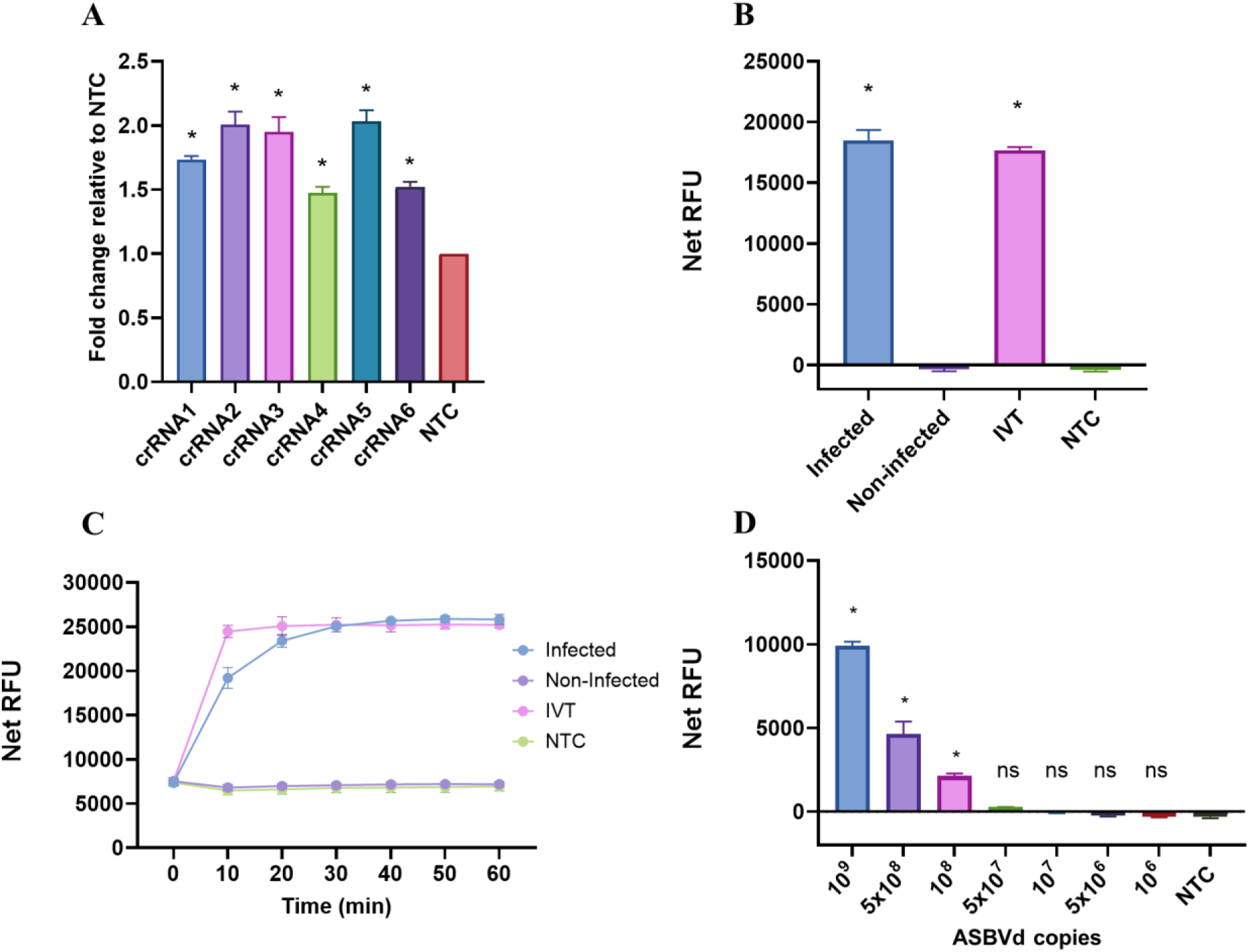
Development and validation of a Cas13a-based detection platform for ASBVd. (A) Screening of six ASBVd-targeting crRNAs using *in vitro*–transcribed (IVT) ASBVd RNA. (B) Detection of ASBVd in RNA extracted from infected avocado tissues using the Cas13a assay. (C) Kinetic fluorescence analysis of ASBVd detection in RNA extracted from infected avocado tissues using the Cas13a assay. (D) Sensitivity evaluation of direct Cas13a-based ASBVd detection using serial dilutions of RNA extracted from ASBVd-infected avocado samples. (A, B and D) *, statistically significant *p* = 0.001; ns, non-significant.

To validate the detection platform with biological samples, RNA extracted from ASBVd-infected avocado tissues was directly subjected to Cas13a detection in 10 min (Figure 6B and Figure 6C). Infected samples generated significantly elevated fluorescence signals compared with non-infected controls, confirming successful detection of ASBVd in real plant-derived RNA samples. Sensitivity analysis using serially diluted infected plant RNA further demonstrated a limit of detection of approximately 10^8^ ASBVd copies in biological samples (Figure 6D), which was lower than the sensitivity observed using *in vitro*-transcribed RNA standards (Supplementary Figure S7). This reduced sensitivity is likely due to the strong inhibitory effect of avocado total RNA on the Cas13a detection reaction (Supplementary Figure S8). Together, these results demonstrate that the developed Cas13a platform can be readily reprogrammed for detection of viroids belonging to distinct taxonomic families through simple crRNA redesign, highlighting its modularity, adaptability, and broad potential for viroid diagnostics.

## Discussion

CRISPR-based nucleic acid detection systems have emerged as attractive alternatives to conventional molecular diagnostics because of their programmability, high specificity, and compatibility with rapid and portable detection formats. For viroid diagnostics, Cas13a offers particular advantages because viroids are RNA pathogens that replicate exclusively through RNA intermediates (Hao et al., 2024; Kalantidis et al., 2025). In contrast, Cas12-based systems generally require RT and target amplification prior to detection, increasing workflow complexity and the risk of amplification-associated false positives. Although Cas13a-based SHERLOCK platforms have previously been applied for viroid detection, these systems still depend on pre-amplification and additional *in vitro* transcription steps before Cas13a activation (Zhai et al., 2024). Several amplification-free Cas13a detection strategies have also been reported; however, many require specialized microfluidic devices (Chandrasekaran et al., 2022; Shan et al., 2023), microchamber-array systems (Shinoda et al., 2021), or electrochemical biosensors (Heo et al., 2022; Qu et al., 2025) that are not readily accessible in standard molecular biology laboratories. In the plant field, amplification-free CRISPR diagnostics remain relatively underexplored, likely due to the complexity of plant-derived samples and the strong inhibitory effects associated with abundant host RNAs. Although several studies have reported CRISPR-based detection of plant viruses and viroids, most still rely on target amplification prior to detection (Besati et al., 2025; Jiao et al., 2021; Marqués et al., 2022; Xu et al., 2026).

This study demonstrates direct amplification-free Cas13a-based detection of plant viroids. Compared with the previously reported SHERLOCK-based viroid detection system, the developed platform eliminated both target amplification and additional *in vitro* transcription steps while maintaining improved analytical sensitivity. crRNAs screening was particularly important because viroid RNAs contain highly stable secondary structures that can restrict crRNA accessibility and Cas13a activation efficiency. Lamb *et al*. (Lamb et al., 2025) have reported that combining multiple crRNAs can further enhance amplification-free Cas13a sensitivity for viral detection. However, under our experimental conditions, multiplexed crRNA combinations did not further improve the detection limit for PSTVd, suggesting that RNA viroid structural constraints and target accessibility may represent important limiting factors in amplification-free viroid detection. Importantly, systematic crRNA screening identified both a highly specific PSTVd-targeting crRNA and a broader-spectrum crRNA capable of detecting multiple pospiviroids. Species-specific detection of viroids remains challenging because of the high sequence conservation among closely related viroid species, which can complicate primer design and reduce assay specificity in conventional RT-PCR-based methods. Our findings further support the potential of Cas13a-based diagnostics to achieve high sequence specificity through optimized crRNA selection while maintaining sensitive RNA detection.

Although phenol/CF11/LiCl workflow produced the strongest signals, the simplified phenol/LiCl workflow maintained robust detection performance while substantially reducing workflow complexity and processing time. Because CF11 cellulose chromatography is increasingly difficult to access commercially, the simplified LiCl-based enrichment strategy and silica-based extraction provide a more practical alternative for broader implementation. The simplified workflows also improve the potential applicability of the platform for field-deployable plant diagnostics. Unlike RT-qPCR, which requires expensive laboratory instrumentation and trained personnel, Cas13a fluorescence outputs can potentially be adapted to portable fluorescence readers, lateral flow strips, or simple UV-based visualization systems. Extension of the platform to ASBVd further demonstrated the adaptability of the detection strategy across phylogenetically distinct viroids. However, ASBVd sample preparation required additional polysaccharide removal steps, likely due to the high polysaccharide content of avocado tissues leading to the stronger effect on Cas13a reaction assay compared to tomato tissue. This observation suggests that optimization of RNA preparation workflows may still be necessary for different plant species and tissue types.

Overall, this study establishes a practical amplification-free Cas13a-based platform for direct viroid detection in plant samples. By combining optimized crRNA selection with simplified RNA enrichment workflows, the developed system achieved sensitive and specific viroid detection in few minutes while reducing assay complexity compared with previously reported amplification-dependent CRISPR diagnostics. These findings provide a foundation for future development of rapid, portable, and field-deployable CRISPR-based detection systems for plant RNA pathogens.

## Materials and Methods

### Plant materials and viroid inoculation

Micro-Tom tomato plants were used for all experiments unless otherwise stated. For robustness evaluation experiments, four tomato varieties, Rutgers, Micro-Tom, Moneymaker, and Marglobe, were used. Plants were inoculated with PSTVd (GenBank accession no. U23058) by agroinfiltration (Bedoya and Daròs, 2010) and maintained under identical greenhouse conditions at 24 °C throughout the experiment. All plants were inoculated at the same developmental stage and sampled at identical time points to minimize experimental variation. Non-inoculated plants were used as negative controls. Unless otherwise stated, leaf samples were collected at 4 weeks post agroinfiltration (wpi). For the time-course experiment (Figure 4B), leaf samples were collected at 1, 2, 3, and 4 wpi. Non-infected and ASBVd (GenBank accession no. X52041)-infected avocado (cultivar Fuerte) trees are maintained at the IBMCP, CSIC-UPV (Valencia, Spain) greenhouse for long time.

### RNA extraction and viroid-enrichment methods

Viroid-enriched RNA was prepared using CF11 cellulose chromatography following the protocol previously described (Pallás et al., 1987). Briefly, plant tissues were homogenized in phenol-based extraction buffer, and aqueous RNA fractions were subjected to CF11 cellulose chromatography for enrichment of low-molecular-weight RNAs. For LiCl-enriched workflows, 4 M LiCl was added at an equal volume after RNA extraction by phenol and/or CF11 enrichment. Samples were incubated to selectively precipitate high-molecular-weight host RNAs, whereas viroid-associated low-molecular-weight RNAs remained in the soluble fraction. The soluble fraction was subsequently recovered by ethanol precipitation. Silica column-based RNA extraction was performed using Zymo-Spin I columns (Zymo Research) following a modified guanidinium thiocyanate-based extraction protocol. Briefly, 0.5 g plant tissue was ground in liquid nitrogen and homogenized in TEX extraction buffer (4 M guanidinium thiocyanate, 0.1 M sodium acetate pH 5.5, 10 mM EDTA, and 0.2 M 2-mercaptoethanol). Following centrifugation, the supernatant was mixed with ethanol and loaded onto Zymo-Spin I columns. Columns were washed twice with TLA buffer (70% ethanol, 10 mM sodium acetate pH 5.5), and RNA was eluted in TEL buffer (20 mM Tris-HCl pH 8.5).

### crRNA design and cloning, and RNA *in vitro* transcription

Cas13a crRNAs targeting PSTVd were designed using the CRISPR RNA design webserver developed by the New York Genome Center (https://cas13design.nygenome.org). Candidate spacer sequences were selected based on BLAST analysis against the *Solanum tuberosum* transcriptome to minimize predicted off-target alignment. Spacer sequences of 28 nucleotides were used for crRNA construction. Spacer sequences were fused to a 5′ direct repeat sequence to generate complete crRNAs and cloned into the pUT7B vector. Plasmid sequences were experimentally confirmed by the Oxford Nanopore technology (Plasmidsaurus). All the sequences used in this work is in Supplementary Table 1. Plasmids were linearized by BsaI digestion prior to *in vitro* transcription using the TranscriptAid T7 High Yield Transcription Kit (Thermo Fisher Scientific) according to the manufacturer’s instructions. Transcribed crRNAs were purified using RNA Clean & Concentrator spin columns (Zymo Research) and quantified using a NanoDrop spectrophotometer. The single-stranded RNA (ssRNA) fluorescent reporter labelled with fluorescein at the 5′ end and a dark quencher at the 3′ end (5′-FAM-UUUUU-Iowa Black-3′) was synthesized by Integrated DNA Technologies (IDT).

### Cas13a detection assay

Cas13a detection assays were performed using the GenCRISPR Cas13a (C2c2) nuclease system (GenScript, Z03486). To assemble the Cas13a/crRNA complex, 45 nM crRNA and 10 nM Cas13a protein were mixed in 1× Cas13a reaction buffer in a total volume of 10 µl and incubated at 37 °C for 10 min. Detection reactions were prepared in a final volume of 20 µl containing 1× Cas13a reaction buffer, 10 pmol fluorescent ssRNA reporter, target RNA sample, nuclease-free water, and 10 µl of preassembled Cas13a/crRNA complex. Reactions were incubated at 37 °C for 60 min unless otherwise stated. Fluorescence signals were measured using a microplate reader (CFX96 real-time system, BioRad) with excitation and emission wavelengths of 494 nm and 518 nm, respectively. For kinetic analyses, fluorescence signals were monitored every 10 min.

## Data analysis

Fluorescence data were analyzed using GraphPad Prism software. Net fluorescence intensity was calculated using the following equation: fluorescence intensity (at the corresponding time point) − fluorescence intensity (at time 0). Data are presented as mean ± standard deviation (SD) from three experimental or biological replicates unless otherwise stated. Statistical analyses were performed as indicated in the corresponding figure legends and included one-way ANOVA followed by Tukey’s or Dunnett’s multiple comparisons tests, two-way ANOVA, and linear regression analysis where appropriate. Statistical significance was defined as p <= 0.05. Asterisks indicate statistically significant differences, whereas “ns” indicates non-significant differences.

## Acknowledgements

This research was supported by the Ministerio de Ciencia, Innovación y Universidades (MCIU; Spain) through the Agencia Estatal de Investigación (PID2021-127671NB-I00, PDC2022-133941-I00 and PID2023-146418OB-I00; MICIU/AEI/10.13039/501100011033 and ERDF, EU), Generalitat Valenciana through program PROMETEO (CIPROM/2022/21) and European Commission (ViroiDoc, 101169421). L.T.T.L was supported by the European Commission MSCA doctoral network ViroiDoc, 101169421.

## SUPPLEMENTARY INFORMATION

## Supplementary Figures

**Supplementary Figure S1.**
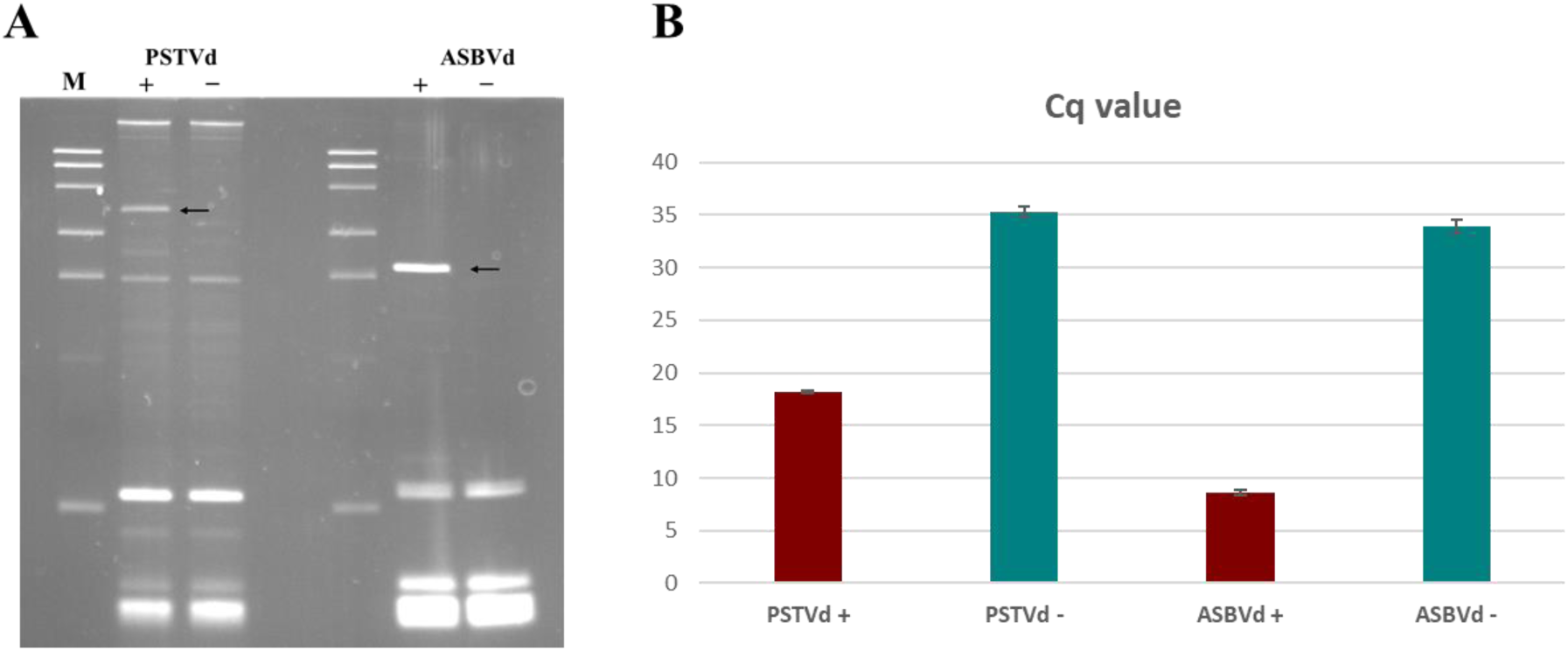
Confirmation of PSTVd- and ASBVd-infected plants by (A) 5% non-denaturing PAGE analysis and (B) RT-qPCR.

**Supplementary Figure S2.**
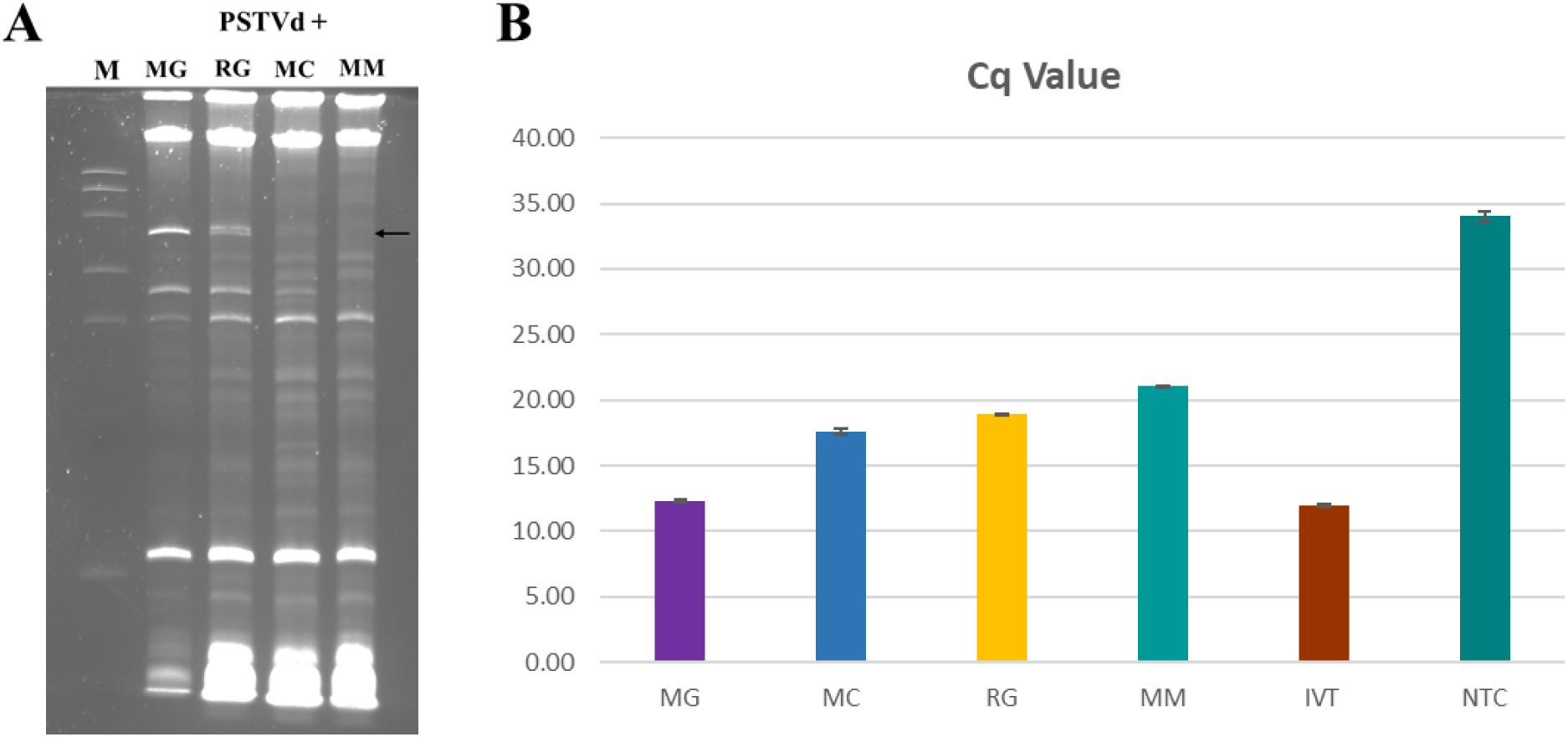
Confirmation of PSTVd infection in four tomato varieties by (A) 5% non-denaturing PAGE analysis and (B) RT-qPCR.

**Supplementary Figure S3.**
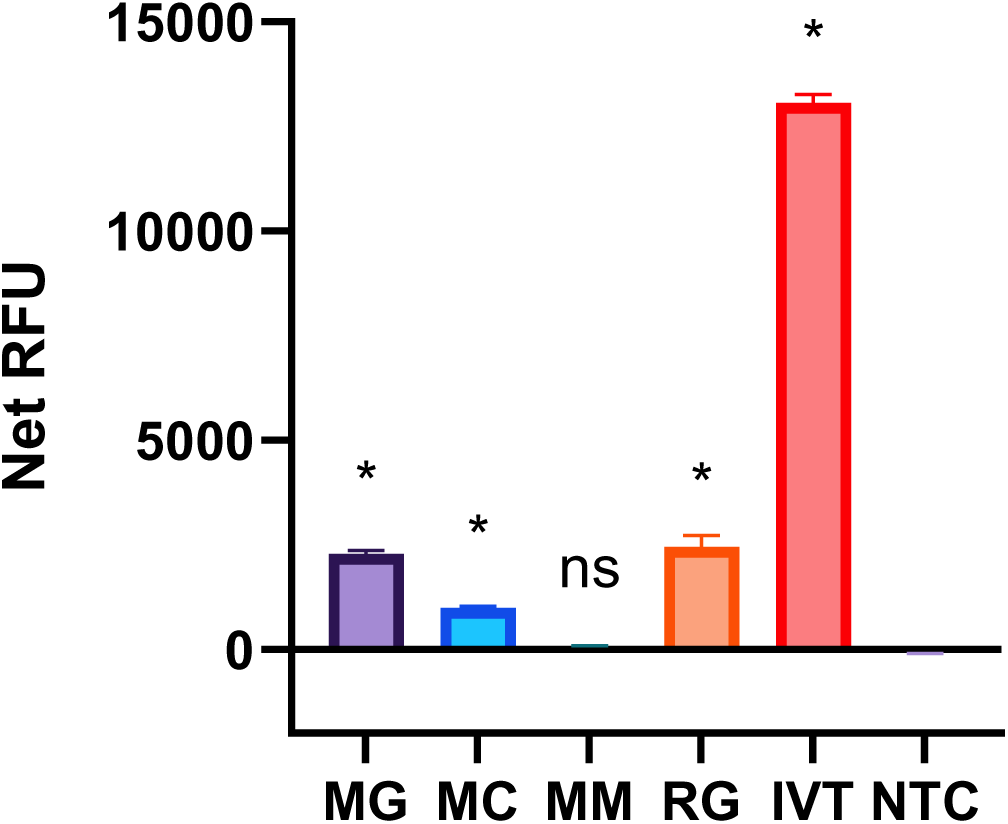
Cas13a-based detection of PSTVd in four tomato varieties using silica column-purified RNA samples.

**Supplementary Figure S4.**
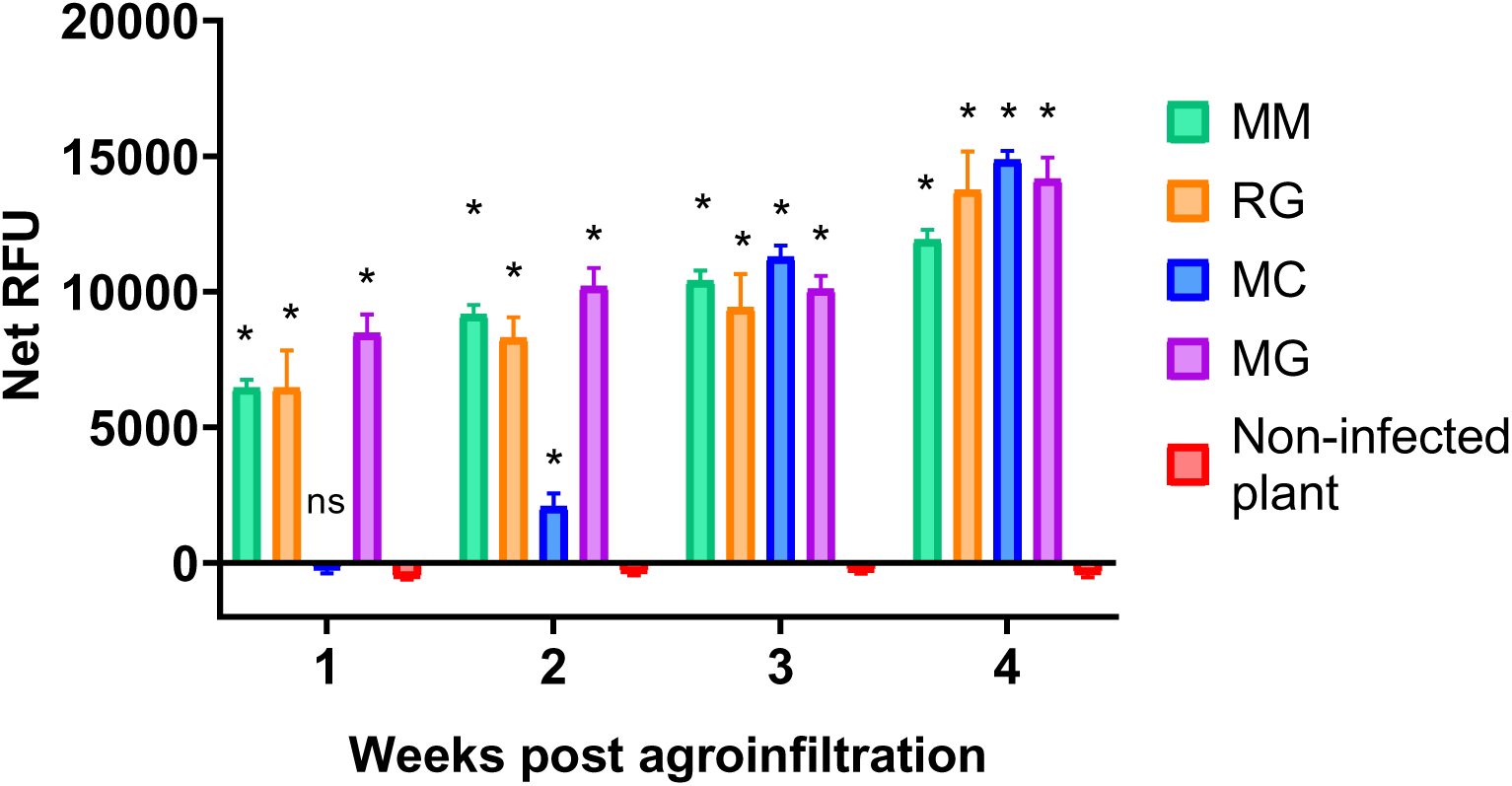
Statistical analysis corresponding to the time-course Cas13a detection of PSTVd shown in Figure 4B.

**Supplementary Figure S5:**
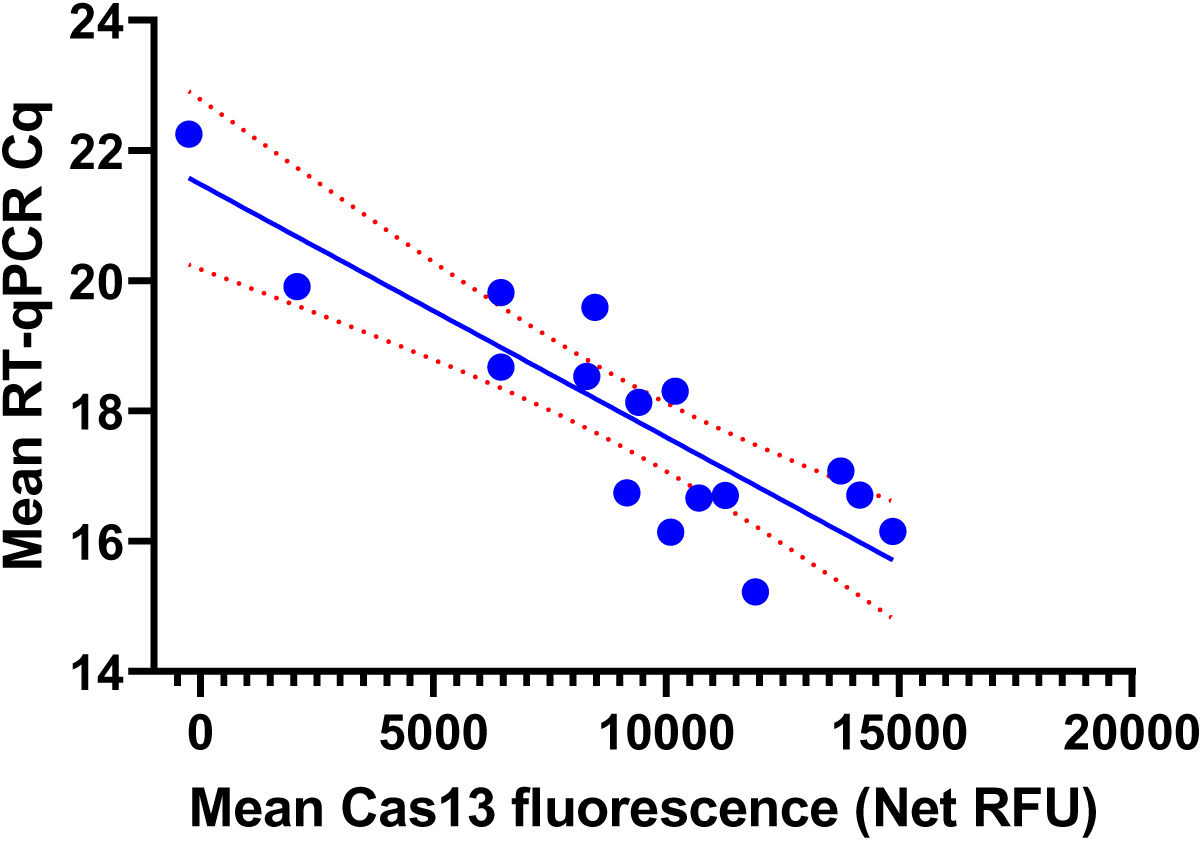
Inverse correlation between Cas13a fluorescence and RT-qPCR Cq values during early PSTVd infection. Scatter plot showing the relationship between Cas13a fluorescence (Net RFU) and RT-qPCR Cq values obtained from four tomato varieties sampled at 1, 2, 3, and 4 weeks post agroinfiltration. Each data point represents the mean of three biological replicates. Pearson correlation analysis revealed a strong inverse correlation between Cas13a fluorescence and RT-qPCR Cq values (*r* = −0.8631, 95% CI = −0.9516 to −0.6421). The solid line represents the linear regression fit, and the dashed lines indicate the 95% confidence interval.

**Supplementary Figure S6.**
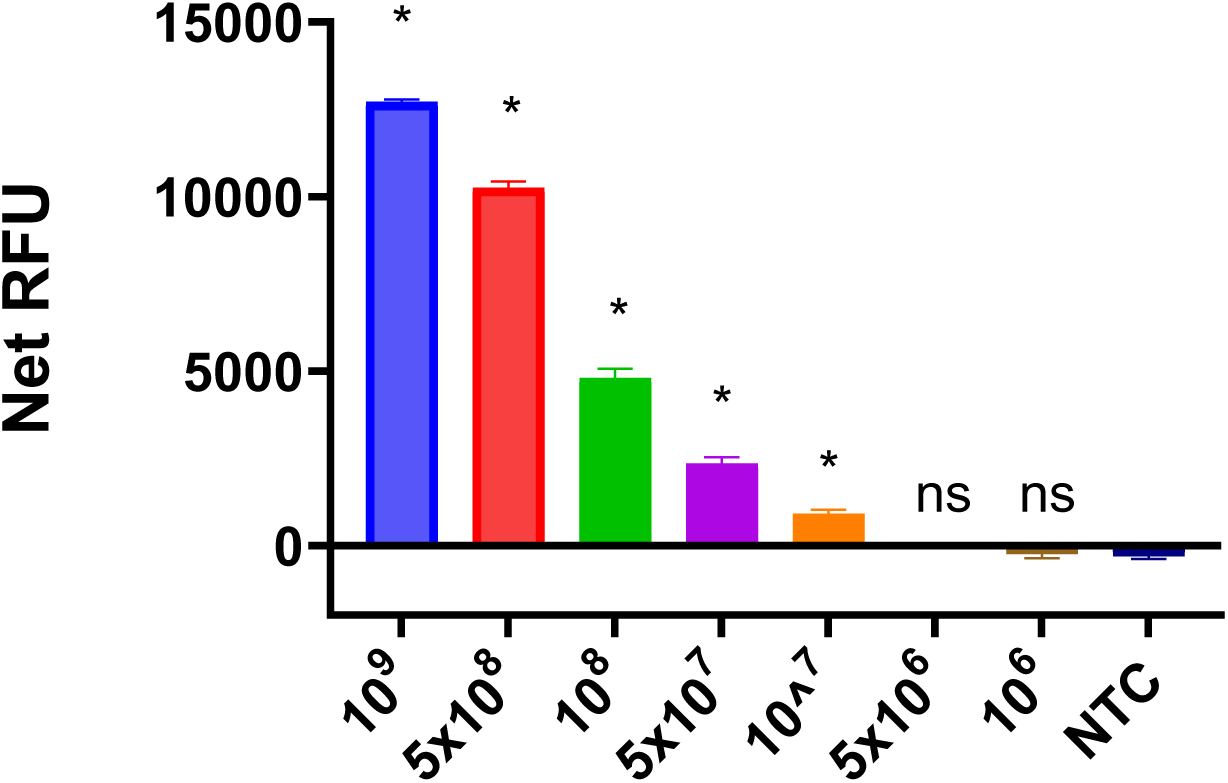
Cas13a detection sensitivity (limit-of-detection) analysis of ASBVd using in vitro–transcribed RNA.

**Supplementary Figure S7.**
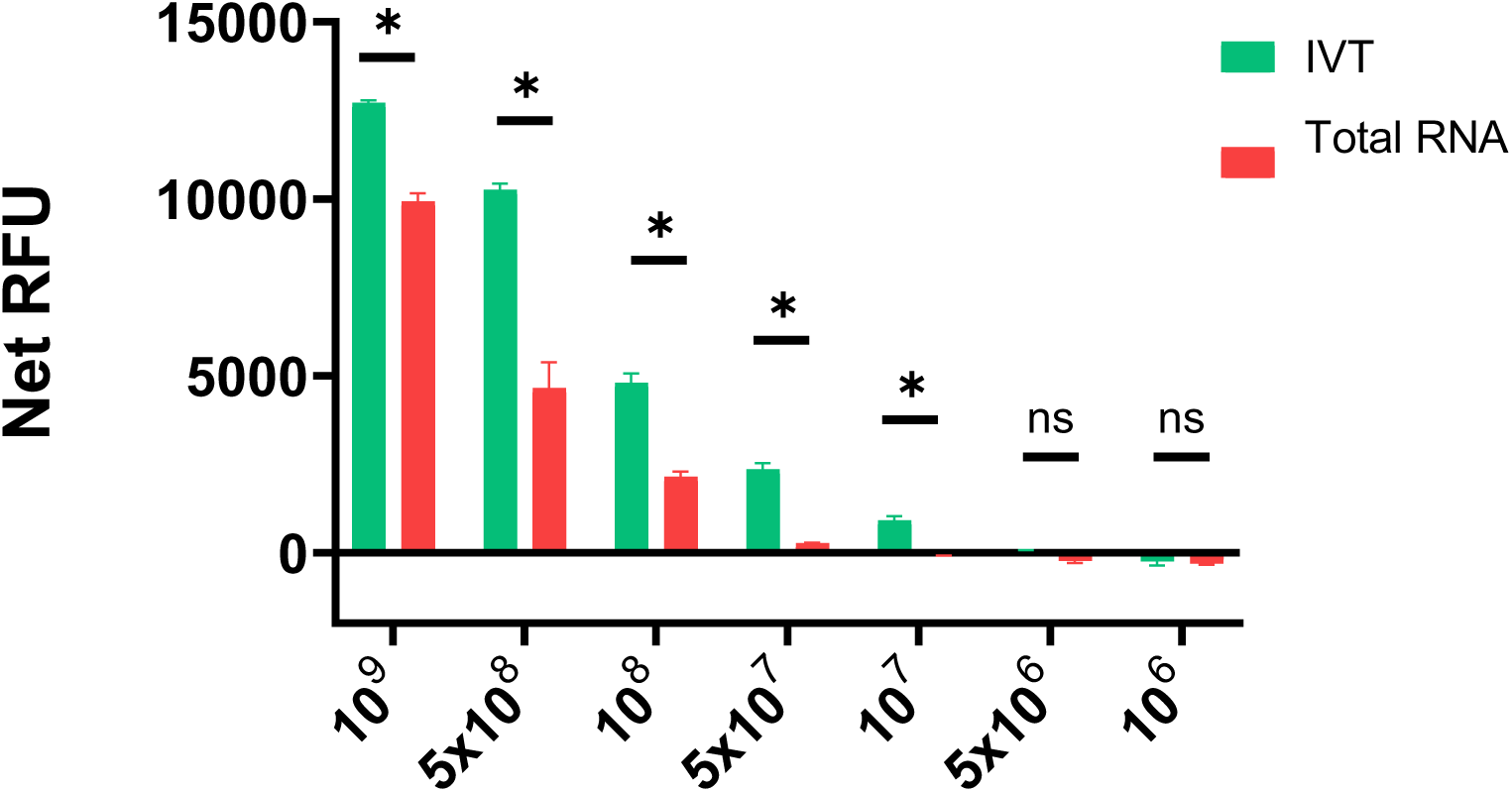
Comparison of Cas13a detection sensitivity (limit of detection) between in vitro–transcribed ASBVd RNA and total RNA extracted from ASBVd-infected plants.

**Supplementary Figure S8.**
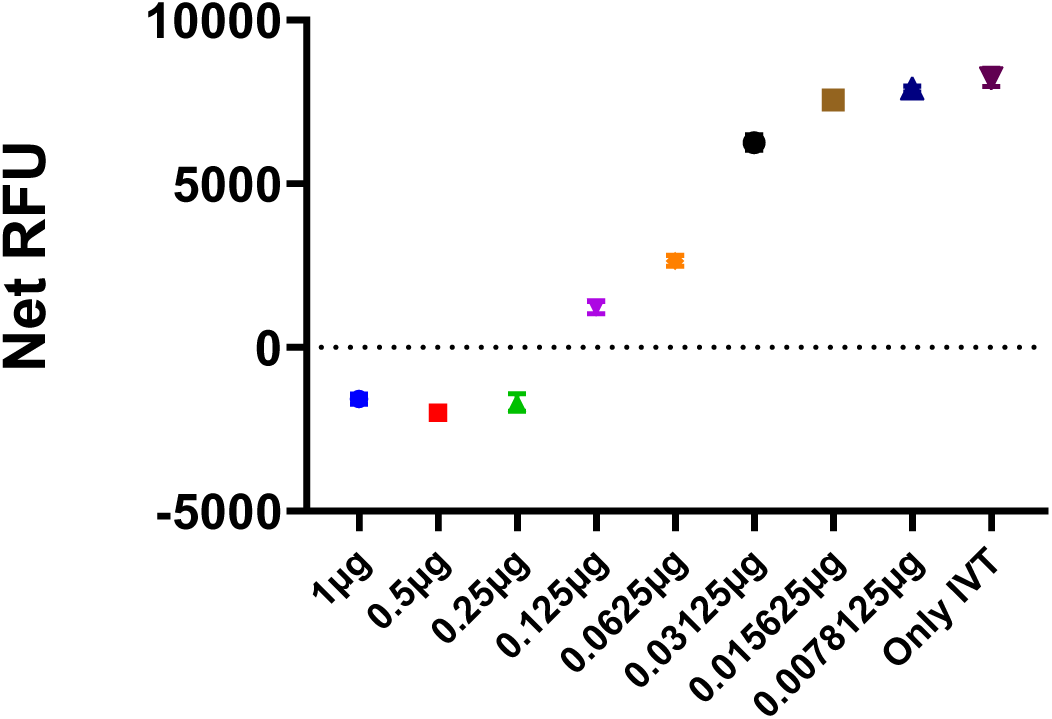
Effect of plant-derived background RNA on the limit of detection of the Cas13a detection assay.

## Supplementary Tables

**Table. S1.**
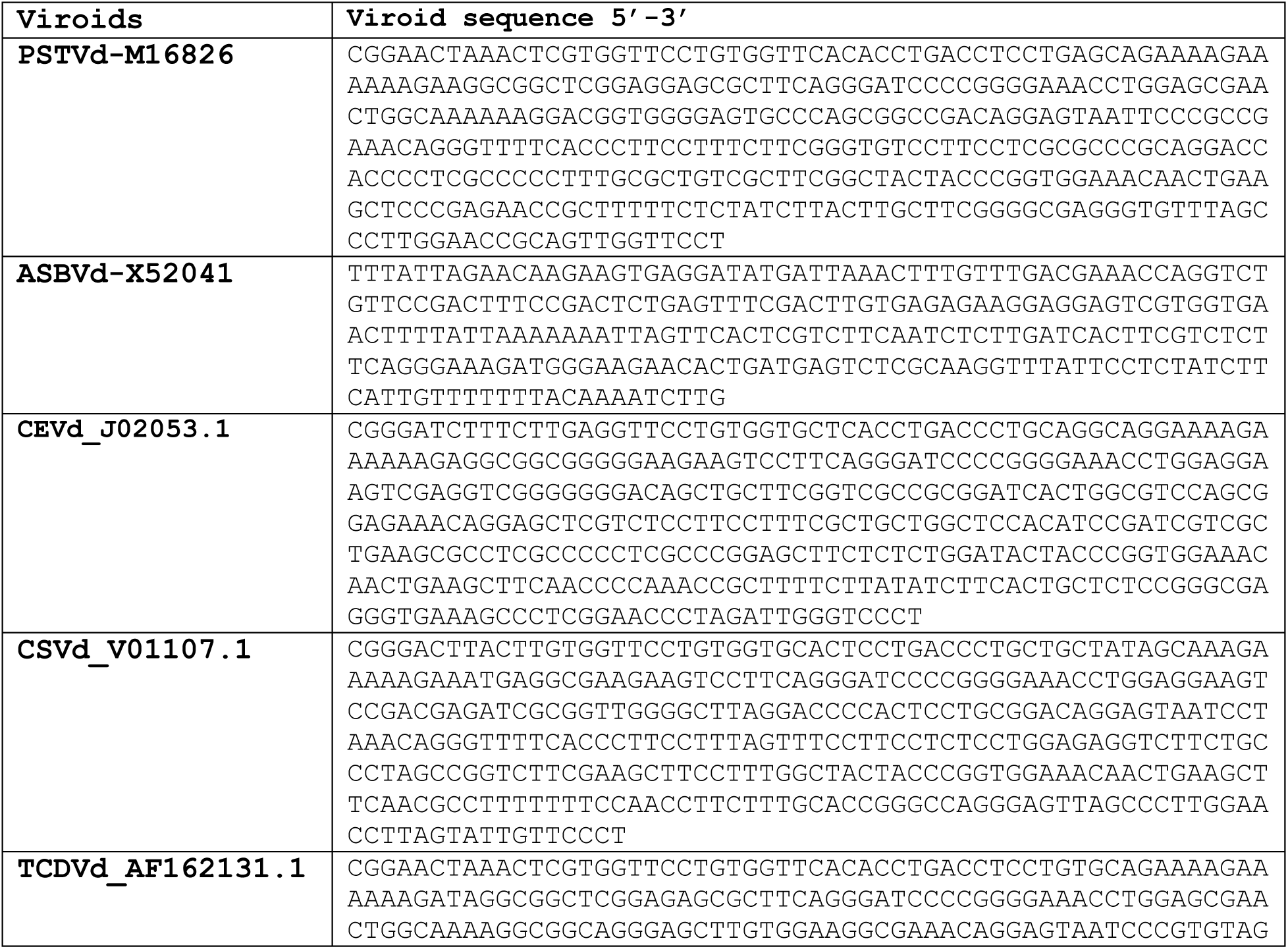
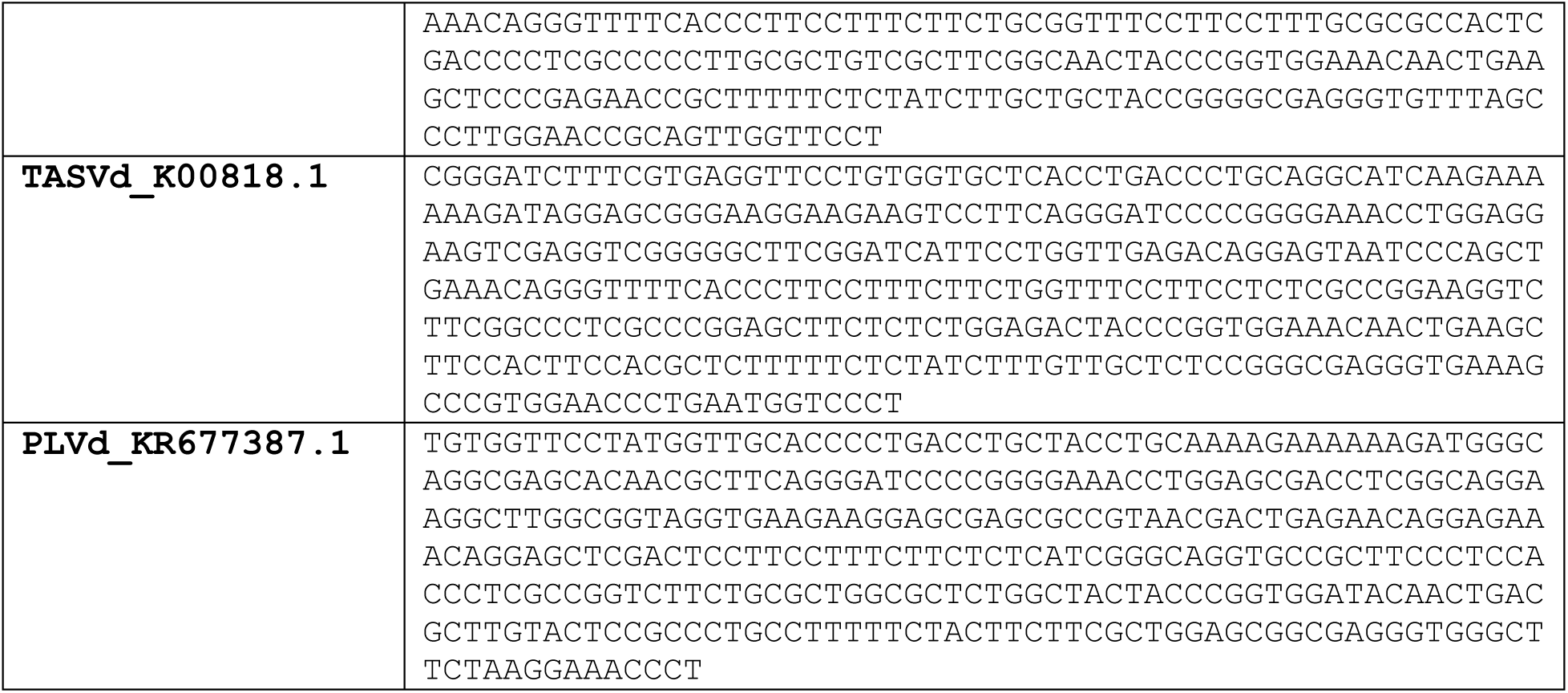
Viroid sequences.

**Table S2.**
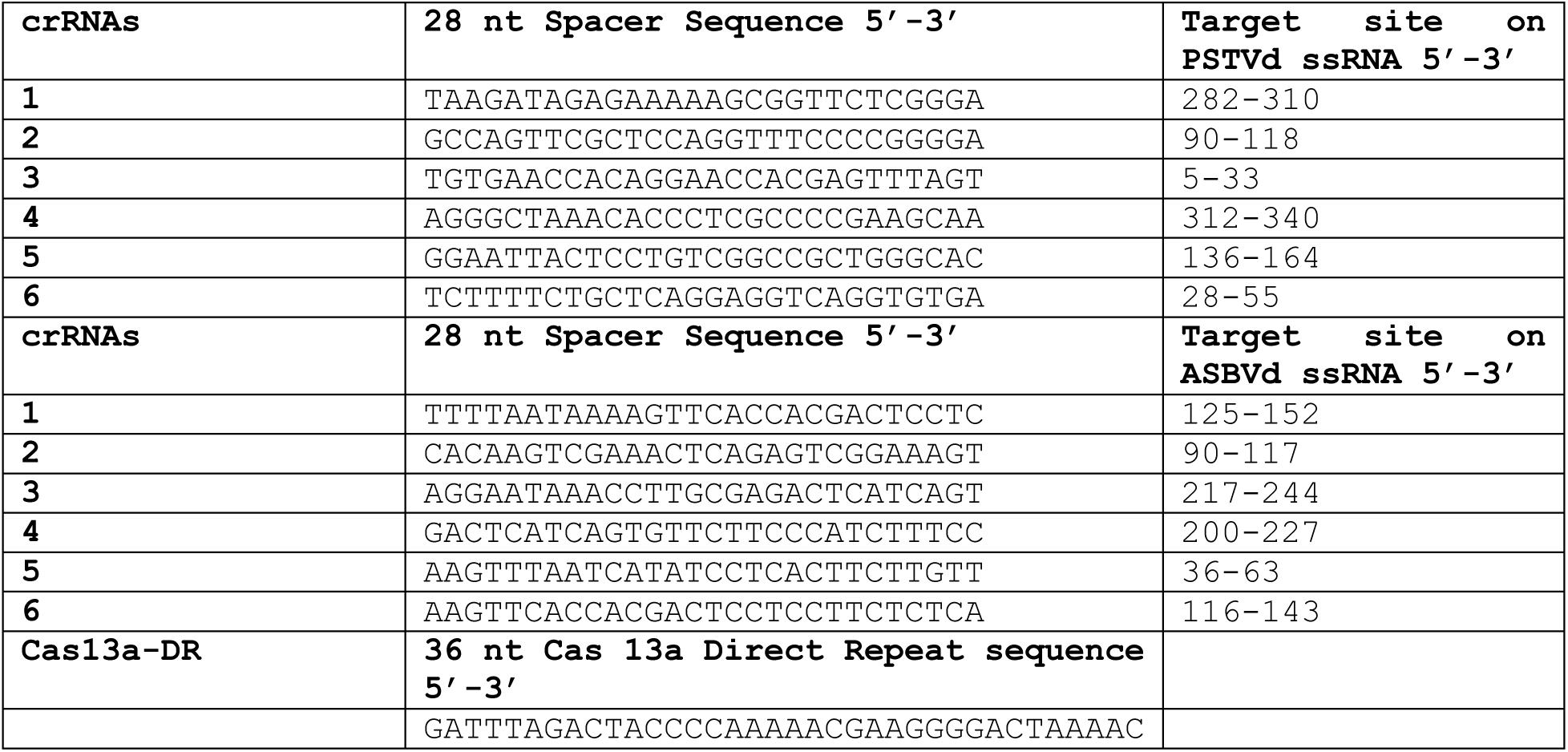
Sequences of crRNAs and Cas13a direct repeat.

**Table S3.**
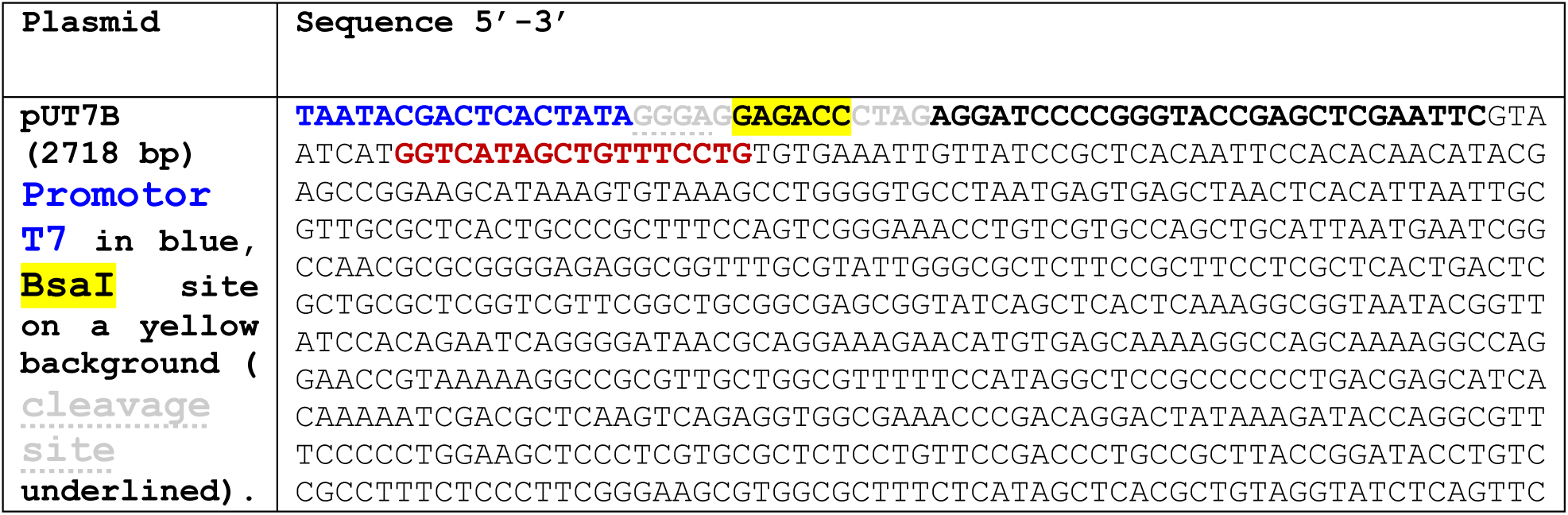
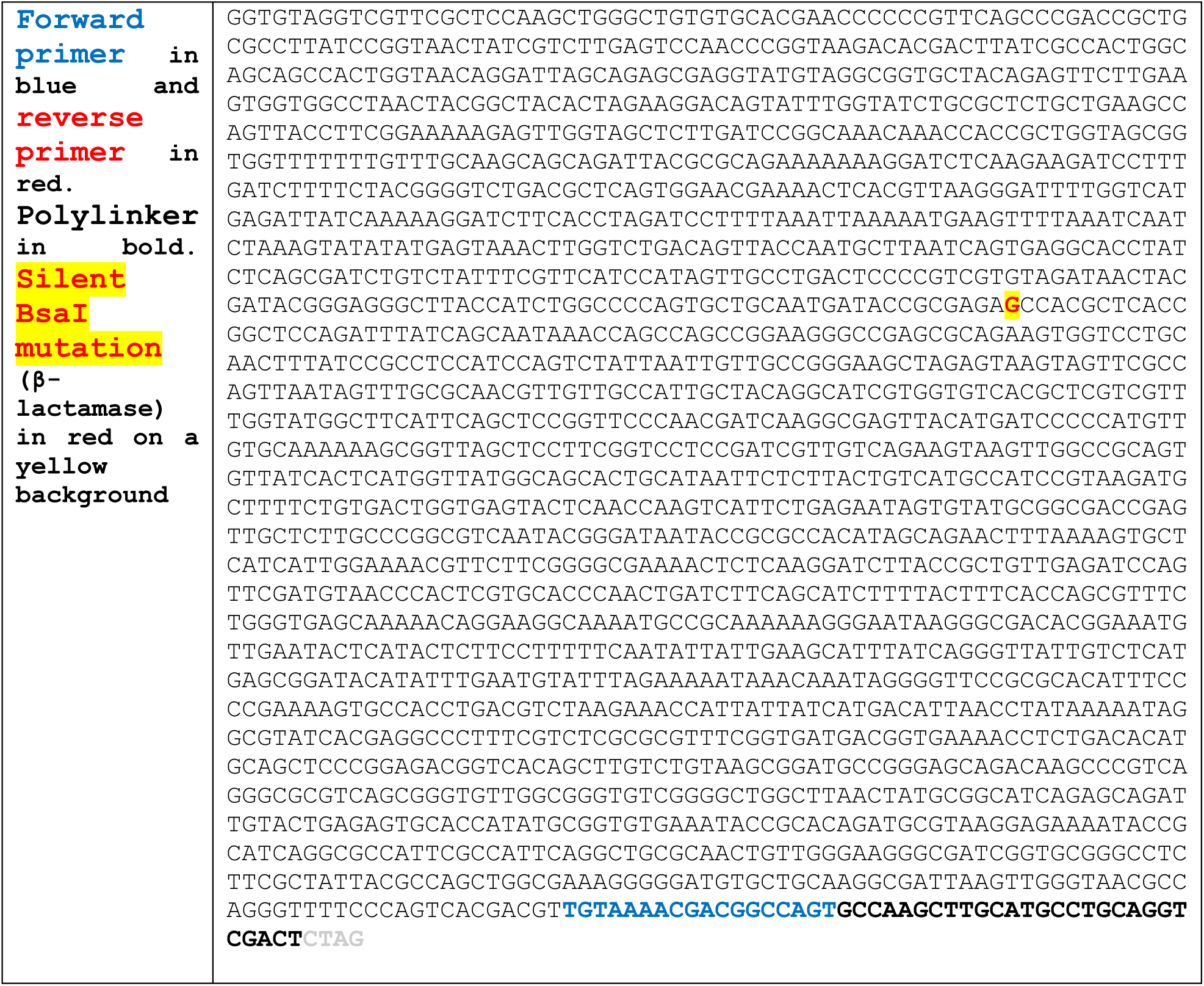
Sequence of pUT7B plasmid.

## Notes

### Competing Interest Statement

The authors have declared no competing interest.

## References

Anwar, S., Khan, S., Azmi, I., Islam, K. U., Ahmad, T., and Iqbal, J. (2025). CRISPR-based molecular detection of SARS-CoV-2, its emerging variants, and diverse pathogens. Diagnostic Microbiology and Infectious Disease 113, 117062.

Bedoya, L. C., and Daròs, J. A. (2010). Stability of Tobacco etch virus infectious clones in plasmid vectors. Virus Res 149, 234–40.

Besati, M., Safarnejad, M. R., Aliahmadi, A., Farzaneh, M., Ruiz, R., Montagud-Martínez, R., Rodrigo, G., and Rafati, H. (2025). Detection of tomato brown rugose fruit virus through CRISPR-Cas12a and CRISPR-Cas9 systems. Sci Rep 15, 25638.

Chandrasekaran, S. S., Agrawal, S., Fanton, A., Jangid, A. R., Charrez, B., Escajeda, A. M., Son, S., McIntosh, R., Tran, H., Bhuiya, A., de León Derby, M. D., Switz, N. A., Armstrong, M., Harris, A. R., Prywes, N., Lukarska, M., Biering, S. B., Smock, D. C. J., Mok, A., Knott, G. J., Dang, Q., Van Dis, E., Dugan, E., Kim, S., Liu, T. Y., Hamilton, J. R., Lin-Shiao, E., Stahl, E. C., Tsuchida, C. A., Giannikopoulos, P., McElroy, M., McDevitt, S., Zur, A., Sylvain, I., Ciling, A., Zhu, M., Williams, C., Baldwin, A., Moehle, E. A., Kogut, K., Eskenazi, B., Harris, E., Stanley, S. A., Lareau, L. F., Tan, M. X., Fletcher, D. A., Doudna, J. A., Savage, D. F., Hsu, P. D., and Consortium, I. G. I. T. (2022). Rapid detection of SARS-CoV-2 RNA in saliva via Cas13. Nature Biomedical Engineering 6, 944–956.

Chertow, D. S. (2018). Next-generation diagnostics with CRISPR. Science 360, 381–382.

Di Serio, F., Owens, R. A., Li, S. F., Matoušek, J., Pallás, V., Randles, J. W., Sano, T., Verhoeven, J. T. J., Vidalakis, G., Flores, R., and Ictv Report, C. (2021). ICTV Virus Taxonomy Profile: Pospiviroidae. J Gen Virol 102.

Di Serio, F., Owens, R. A., Navarro, B., Serra, P., Martínez de Alba Á, E., Delgado, S., Carbonell, A., and Gago-Zachert, S. (2023). Role of RNA silencing in plant-viroid interactions and in viroid pathogenesis. Virus Res 323, 198964.

Diener, T. O. (1971). Potato spindle tuber “virus”: IV. A replicating, low molecular weight RNA. Virology 45, 411–428.

Gong, X.-Y., Wang, Z.-H., Bashir, M., Tang, T., Gan, X., and Yang, W.-C. (2025). Recent application of CRISPR/Cas in plant disease detection. TrAC Trends in Analytical Chemistry 189, 118251.

Gootenberg, J. S., Abudayyeh, O. O., Lee, J. W., Essletzbichler, P., Dy, A. J., Joung, J., Verdine, V., Donghia, N., Daringer, N. M., Freije, C. A., Myhrvold, C., Bhattacharyya, R. P., Livny, J., Regev, A., Koonin, E. V., Hung, D. T., Sabeti, P. C., Collins, J. J., and Zhang, F. (2017). Nucleic acid detection with CRISPR-Cas13a/C2c2. Science 356, 438–442.

Guček, T., Jakše, J., and Radišek, S. (2023). Optimization and Validation of Singleplex and Multiplex RT-qPCR for Detection of Citrus bark cracking viroid (CBCVd), Hop latent viroid (HLVd), and Hop stunt viroid (HSVd) in Hops (Humulus lupulus). Plant Dis 107, 3592–3601.

Hajeri, S., Vidalakis, G., and Yokomi, R. K. (2022). Detection of Viroids Using RT-qPCR. Methods Mol Biol 2316, 153–162.

Hammond, R. W. (2017). Chapter 1 - Economic Significance of Viroids in Vegetable and Field Crops. *In* “Viroids and Satellites” (A. Hadidi, R. Flores, J. W. Randles and P. Palukaitis, eds.), pp. 5–13. Academic Press, Boston.

Hao, J., Ma, J., and Wang, Y. (2024). Understanding viroids, endogenous circular RNAs, and viroid-like RNAs in the context of biogenesis. PLoS Pathog 20, e1012299.

Heo, W., Lee, K., Park, S., Hyun, K. A., and Jung, H. I. (2022). Electrochemical biosensor for nucleic acid amplification-free and sensitive detection of severe acute respiratory syndrome coronavirus 2 (SARS-CoV-2) RNA via CRISPR/Cas13a trans-cleavage reaction. Biosens Bioelectron 201, 113960.

Hussain, W., Li, G., Ain, Q. U., Spetz, C., Lv, D., and Zhou, Y. (2026). Potato Spindle Tuber Viroid (PSTVd)-Plant Interactions: Molecular Mechanisms, Multi-Omics Reprogramming, and Emerging Ecological Dynamics of the Holobiont. Plant Cell Environ.

Jaybhaye, S. G., Chavhan, R. L., Hinge, V. R., Deshmukh, A. S., and Kadam, U. S. (2024). CRISPR-Cas assisted diagnostics of plant viruses and challenges. Virology 597, 110160.

Jiao, J., Kong, K., Han, J., Song, S., Bai, T., Song, C., Wang, M., Yan, Z., Zhang, H., Zhang, R., Feng, J., and Zheng, X. (2021). Field detection of multiple RNA viruses/viroids in apple using a CRISPR/Cas12a-based visual assay. Plant Biotechnol J 19, 394–405.

Kalantidis, K., Tselika, M., Kallemi, P., Bardani, E., Kryovrysanaki, N., and Katsarou, K. (2025). Derailing the host machinery to achieve replication: how viroid and viroid-like RNAs successfully copy their genomes in hostile territory. RNA Biol 22, 1–19.

Khan, W. A., Barney, R. E., and Tsongalis, G. J. (2021). CRISPR-cas13 enzymology rapidly detects SARS-CoV-2 fragments in a clinical setting. J Clin Virol 145, 105019.

Lamb, C. H., Riesle-Sbarbaro, S., Prescott, J. B., Te Velthuis, A. J. W., Myhrvold, C., and Nilsson-Payant, B. E. (2025). Amplification-free detection of zoonotic viruses using Cas13 and multiple CRISPR RNAs. J Gen Virol 106.

Leichtfried, T., Reisenzein, H., Steinkellner, S., and Gottsberger, R. A. (2021). Improvement in the sensitivity of viroid detection by adapting the reverse transcription step in one-step RT-qPCR assays. Journal of Virological Methods 292, 114123.

Li, S.-Y., Cheng, Q.-X., Liu, J.-K., Nie, X.-Q., Zhao, G.-P., and Wang, J. (2018). CRISPR-Cas12a has both cis-and trans-cleavage activities on single-stranded DNA. Cell Research 28, 491–493.

Marqués, M. C., Sánchez-Vicente, J., Ruiz, R., Montagud-Martínez, R., and Márquez-Costa, R. (2022). Diagnostics of Infections Produced by the Plant Viruses TMV, TEV, and PVX with CRISPR-Cas12 and CRISPR-Cas13. ACS Synthetic Biology 11, 2384–2393.

Montagud-Martínez, R., and Márquez-Costa, R. (2025). Virus Detection by CRISPR-Cas9-Mediated Strand Displacement in a Lateral Flow Assay. ACS Appl Bio Mater 8, 4221–4229.

Nie, X., and Singh, R. P. (2017). Chapter 33 - Viroid Detection and Identification by Bioassay. *In* “Viroids and Satellites” (A. Hadidi, R. Flores, J. W. Randles and P. Palukaitis, eds.), pp. 347–356. Academic Press, Boston.

Ortolá, B., and Daròs, J. A. (2023). Viroids: Non-Coding Circular RNAs Able to Autonomously Replicate and Infect Higher Plants. Biology (Basel) 12.

Pallás, V., Navarro, A., and Flores, R. (1987). Isolation of a Viroid-like RNA from Hop Different from Hop Stunt Viroid. Journal of General Virology 68, 3201–3205.

Pallás, V., Sánchez-Navarro, J. A., and James, D. (2018). Recent Advances on the Multiplex Molecular Detection of Plant Viruses and Viroids. Front Microbiol 9, 2087.

Pedrelli, A., and Vergine, M. (2026). Inside the European Plant Viroid Scenario: Continental Distribution, Host Range, and Genetic Features of the Main Viroid Populations. Viruses 18.

Qu, L., Meng, L., Sun, X., Cui, W., Shi, J., Wang, Y., Hu, Q., Xu, S., Sun, B., and Liang, C. (2025). CRISPR/Cas-Based Electrochemical Biosensor for Human Immunodeficiency Virus-1 Nucleic Acid Amplification-Free Detection to the Attomolar Level. ACS Sensors 10, 5613–5622.

Sanger, H. L., Klotz, G., Riesner, D., Gross, H. J., and Kleinschmidt, A. K. (1976). Viroids are single-stranded covalently closed circular RNA molecules existing as highly base-paired rod-like structures. Proc Natl Acad Sci U S A 73, 3852–6.

Sano, T. (2021). Progress in 50 years of viroid research-Molecular structure, pathogenicity, and host adaptation. Proc Jpn Acad Ser B Phys Biol Sci 97, 371–401.

Shan, X., Gong, F., Yang, Y., Qian, J., Tan, Z., Tian, S., He, Z., and Ji, X. (2023). Nucleic Acid Amplification-Free Digital Detection Method for SARS-CoV-2 RNA Based on Droplet Microfluidics and CRISPR–Cas13a. Analytical Chemistry 95, 16489–16495.

Shinoda, H., Taguchi, Y., Nakagawa, R., Makino, A., Okazaki, S., Nakano, M., Muramoto, Y., Takahashi, C., Takahashi, I., Ando, J., Noda, T., Nureki, O., Nishimasu, H., and Watanabe, R. (2021). Amplification-free RNA detection with CRISPR–Cas13. Communications Biology 4, 476.

Singh, R. P. (1973). Experimental host range of the potato spindle tuber ‘virus’. American Potato Journal 50, 111–123.

Steger, G., and Perreault, J. P. (2016). Structure and Associated Biological Functions of Viroids. Adv Virus Res 94, 141–72.

Tian, Y., Li, C., Zhao, E., Chen, Y., Shen, X., Xu, S., Yu, Y., and Sun, L. (2026). Recent Advances in the Detection of Plant Diseases Based on the CRISPR-Cas System. Analytical Chemistry 98, 13951–13966.

Venkataraman, S., Shahgolzari, M., Hefferon, K., Atri, E., and De Steur, H. (2024). Economic Impacts of Viroids. Preprints, 10.20944/preprints202405.2134.v1.

Verhoeven, J. T. J. (2010). Identification and epidemiology of pospiviroids. PhD thesis, Wageningen University, Wageningen, The Netherlands.

Xu, J., Jiang, X., Dashtarzhaneh, M. K., Zhong, Y., Sharma, B., Peng, R., Khodadadi, F., Du, K., and Duan, C. (2026). Ultrasensitive, Low-Input Detection of Avocado Sunblotch Viroid via RPA-CRISPR and Nanopore-Array Single-Bead Fluorescence Readout. bioRxiv, 2026.01.19.700023.

Yao, R., Xu, Y., Wang, L., Wang, D., Ren, L., Ren, C., Li, C., Li, X., Ni, W., He, Y., Hu, R., Guo, T., Li, Y., Li, L., Wang, X., and Hu, S. (2021). CRISPR-Cas13a-Based Detection for Bovine Viral Diarrhea Virus. Front Vet Sci 8, 603919.

Zhai, Y., Gnanasekaran, P., and Pappu, H. R. (2024). Development of a CRISPR/SHERLOCK-Based Method for Rapid and Sensitive Detection of Selected Pospiviroids. In “Viruses”, Vol. 16, pp. 1079.

Zhang, Z., and Li, S. (2024). Chapter 16 - Detection of viroids. *In* “Fundamentals of Viroid Biology” (C. R. Adkar-Purushothama, T. Sano, J.-P. Perreault, S. M. Yanjarappa, F. Di Serio and J.-A. Daròs, eds.), pp. 297–321. Academic Press.

Zhou, J., Li, Z., Seun Olajide, J., and Wang, G. (2024). CRISPR/Cas-based nucleic acid detection strategies: Trends and challenges. Heliyon 10, e26179.

